# Having “multiple selves” helps learning agents explore and adapt in complex changing worlds

**DOI:** 10.1101/2022.12.16.520795

**Authors:** Zack Dulberg, Rachit Dubey, Isabel M. Berwian, Jonathan Cohen

## Abstract

Satisfying a variety of conflicting needs in a changing environment is a fundamental challenge for any adaptive agent. Here, we show that designing an agent in a modular fashion as a collection of subagents, each dedicated to a separate need, powerfully enhanced the agent’s capacity to satisfy its overall needs. We used the formalism of deep reinforcement learning to investigate a biologically relevant multi-objective task: continually maintaining homeostasis of a set of physiologic variables. We then conducted simulations in a variety of environments and compared how modular agents performed relative to standard monolithic agents (i.e., agents that aimed to satisfy all needs in an integrated manner using a single aggregate measure of success). Simulations revealed that modular agents: a) exhibited a form of exploration that was intrinsic and emergent rather than extrinsically imposed; b) were robust to changes in non-stationary environments, and c) scaled gracefully in their ability to maintain homeostasis as the number of conflicting objectives increased. Supporting analysis suggested that the robustness to changing environments and increasing numbers of needs were due to intrinsic exploration and efficiency of representation afforded by the modular architecture. These results suggest that the normative principles by which agents have adapted to complex changing environments may also explain why humans have long been described as consisting of ‘multiple selves’.

**Significance Statement:** Adaptive agents must continually satisfy a range of distinct and possibly conflicting needs. In most models of learning, a monolithic agent tries to maximize one value that measures how well it balances its needs. However, this task is difficult when the world is changing and needs are many. Here, we considered an agent as a collection of modules each dedicated to a particular need and competing for control of action. Compared to the standard monolithic approach, modular agents were much better at maintaining homeostasis of a set of internal variables in simulated environments, both static and changing. These results suggest that having ‘multiple selves’ may represent an evolved solution to the universal problem of balancing multiple needs in changing environments.

---

**O**ne of the most fundamental questions about agency is how an individual manages conflicting needs. The question pervades mythology and literature (1, 2), and is a focus of theoretical and empirical work in virtually every scientific discipline that studies agentic behavior, from neuroscience (3), psychology (4–6), economics (7–9) and sociology (10, 11) to artificial intelligence and machine learning (12, 13). Perhaps most famously, the question of how an individual manages conflict has been at the heart of over a century of work on the nature of psychic conflict underlying mental illness (14, 15). How is it that we (and other natural agents) are so effective in managing fluctuating, ongoing and frequently conflicting needs for sustenance, shelter, social interaction, reproduction, temperature regulation, information gathering, etc.? Growing interest in the design of autonomous artificial agents faces similar questions, such as how to balance execution of actions, with the replenishment of energy or need for repair (see Figure 1a). This challenge is especially difficult in a world that is constantly changing (i.e., features of the environment are non-stationary) and when the set of distinct needs is large.

**Fig. 1.**
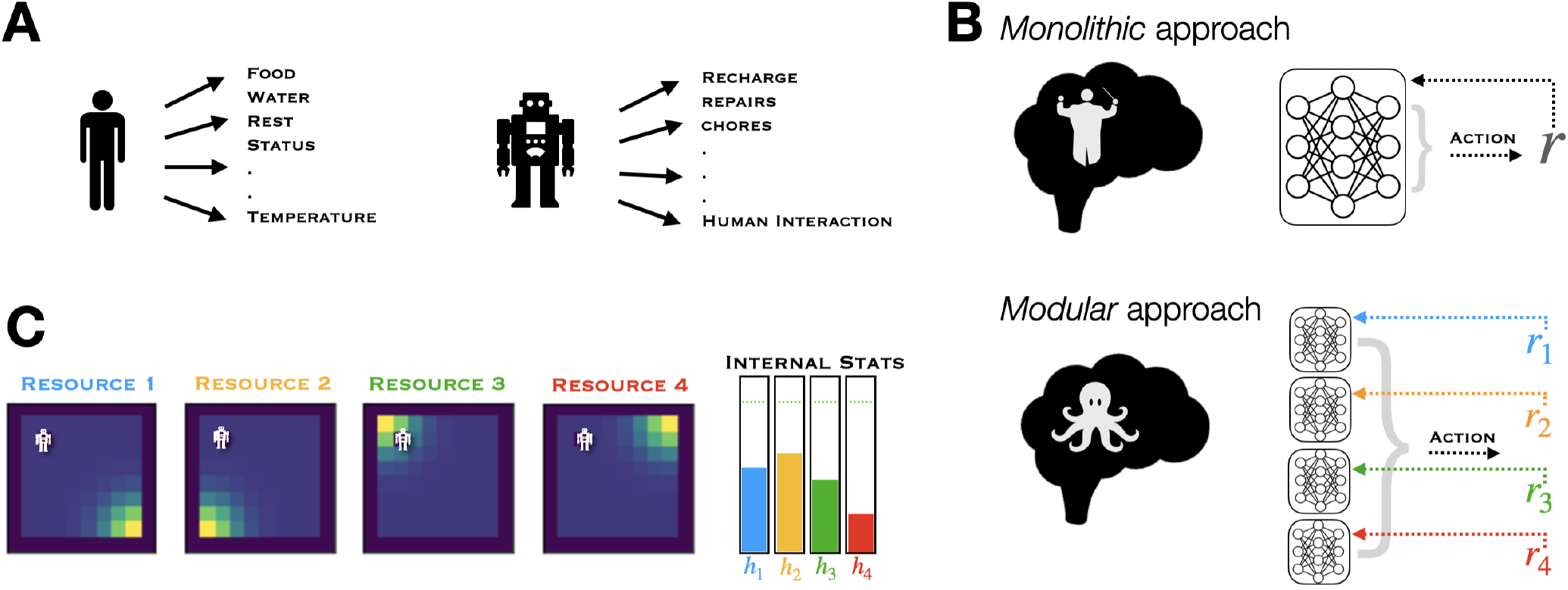
Concept illustrations. **a**: Adaptive organisms are pulled in multiple directions due to competing demands. **b**: Do brains balance our needs in a global fashion (top; *monolithic*) or, like the semi-autonomous legs of an octopus, might sub-agents compete for control? (bottom; *modular* ). In the framework of reinforcement learning, this contrast corresponds to the one between a network learning a single policy based on a scalar reward value *r* (top, right), versus multiple sub-networks each learning a distinct policy based on separate reward components (*r*_1_,*r*_2_,…) (bottom, right). **c**: A homeostatic task environment. An agent (white) moves around a grid-world, searching for densities (yellow) of different resources that can replenish its internal stats (*h*_1_,*h*_2_,…). A distinct resource map is displayed for the distribution of each of the four resources in the same grid-world. Stat “meters” at the right show an example of the levels for each stat at a given point in time, with dotted green lines indicating their set-points.

A central and recurring debate, that arises with the question of how multiple, potentially conflicting needs are managed, is whether this relies on a single, *monolithic* agent (or “self”) that takes integrated account of all needs, or rather reflects an emergent process of competition among multiple *modular* agents (i.e., “multiple selves”) (16–22). In principle, a monolithic system should be capable of translating information about its environment and objectives into intelligent behavior in an integrated fashion. However, the reason modular organization may be so prevalent in biological and psychological systems is because it affords certain benefits in practice.

In this article, we use the computational framework of model-free reinforcement learning (RL) (23) to provide a normative perspective on this debate. We implement two types of agents that must learn to manage multiple needs (or “objectives”): one that treats the problem *monolithically*, as a simultaneous global optimization over the different objectives, and one in which behavior emerges out of competition between sub-agents, each dedicated to a particular objective (Figure 1b). We cast the problem of multiple objectives as the ubiquitous need to maintain homeostatic balance along different dimensions (Figure 1c), and study non-stationarity by introducing changes in the external location of required resources over time. Then, by training deep RL agents in systematically controlled simulated environments, we provide insights about when and why the modular approach, implementing ‘multiple selves’, meets the challenge of learning to manage multiple needs more efficiently and effectively.

The monolithic agent is based on standard principles and mechanisms of RL (23). In this approach, reward is defined as a single scalar value that the agent receives in response to taking actions in its environment. When there are multiple objectives, a scalar reward associated with each is combined into a single reward that the monolithic agent seeks to optimize (13). As a central example, in homeostatically-regulated reinforcement learning (HRRL) (24), an agent with separate homeostatic drives is rewarded based on its ability to maintain all its “homeostats” at their set points. This is done by combining deviations from *all* set-points into a single reward which, when maximized, minimizes homeostatic deviations overall. Not restricted just to primitive drives, HRRL is a general framework for deriving reward from any objective describable by a set-point, whether concrete (like hydration) or abstract (like reaching a goal) (25).

While standard RL is a provably optimal solution for reward maximization in stationary environments, nevertheless this approach faces several challenges. First, an agent must balance collecting known sources of reward with exploring its environment to learn about unknown sources (known as the explore-exploit dilemma (26)), and this typically requires careful tuning of exploratory noise or bonuses (27). Second, standard RL struggles in non-stationary environments (28). Third, it is well known to suffer from the “curse of dimensionality,” which refers to the exponential growth in the number of relationships between states and objectives that must be learned by the agent as the complexity of the environment and/or the number of its objectives increases (29).

In contrast to the monolithic approach, an agent in the modular approach is comprised of separate ‘specialist’ RL sub-agents (modules), each of which learns to optimize reward pertaining to a single need. As a result, different modules learn different policies (action preferences for individual states) and, for a given state, are likely to have different preferences for actions. The action of the agent is selected based on an arbitration of the individual module preferences. Such modular architectures have received increasing attention in RL (30–37), and so has modularity in machine learning more broadly, because this ‘divide-and-conquer’ strategy tends to improve learning speed, scaling properties, and performance robustness (38). However, these approaches typically attempt to decompose a standard scalar reward into a more tractable set of sub-problems. In contrast, our starting point is that ecologically-valid agents have a set of distinct, pre-defined objectives, which can be independent of one another, and the actions needed to meet these may come into conflict. We then ask the question: should these objectives be composed together at the level of reward (monolithic), or at the level of action (modular)? In multi-objective cases like these, dedicating an independent sub-agent to each objective may not be an optimal strategy in all cases (39). However, our hypothesis is that such a one-agent-per-objective heuristic, in which action values rather than rewards are combined, affords practical benefits for an agent that must satisfy independent drives.

Here, we compare modular and monolithic architectures with respect to the ecologically relevant problem of learning to continuously balance multiple homeostatic needs in a changing environment. We find that the modular architecture outperforms the monolithic one, and identify critical factors that contribute to this success by addressing the three aforementioned challenges faced by the monolithic approach. First, we find that exploration emerges naturally as an intrinsic property of the modular architecture, rather than having to be imposed or regulated as an extrinsic factor: modules are often “dragged along” by other modules that currently have the “upper hand” on action, allowing the former to discover the value of actions they themselves would not have selected. Second, by learning to bias their policies toward features that are most relevant to their individual objectives, while ignoring irrelevant features, they are able to adapt effectively to changes in the environment. Together, these factors also make modular agents more robust to an increase in the number of objectives (and therefore the complexity of the task/environment).

In the following sections, we describe the simulations on which the observations summarized above were based. First, we introduce a simple but flexible environment for homeostatic tasks, construct a deep learning implementation of HRRL to learn such tasks, and incorporate this into both monolithic and modular architectures. We then quantify the differences between monolithic and modular agents in the homeostatic tasks. We focus first on an environment with stationary resources, and show that the performance benefits of the modular agent are related to its intrinsic capacities for exploration and efficiency of representation. We then turn to non-stationary environments, and show that as the number of homeostatic needs increases, the relative advantage of modularity is strongly enhanced. Finally, we situate our work within the relevant reinforcement learning, evolutionary biology, neuroscience, and psychology literatures, and suggest that the need to balance multiple objectives in changing environments pressured the evolution of distinct value systems in the brain, providing a normative account for why psychic conflict appears central to human psychology.

## Results

To compare monolithic and modular approaches, we trained deep reinforcement learning agents in a simple grid-world environment. In this environment, various resources were distributed at different locations in the grid. The goal of the agent was to move around the grid to collect resources in order to maintain a set of depleting internal variables (*stats*) at a homeostatic set-point. Stats could be replenished by the visiting the location of the corresponding resource, and could be exceeded through “overconsumption” by remaining too long at that location. For example, if the agent had a low ‘hunger’ stat, it could collect the ‘food’ resource by moving to the location of that resource. If it ate too much (stayed too long), it could leave the location with food until its ‘hunger’ stat decreased below its set point. The monolithic agent was a standard deep Q-network (DQN) (40), while the modular agent was a set of DQNs, one corresponding to each needed resource, from which actions were selected by simply summing up action value outputs across the DQN modules. A more detailed description of the environment and agent design is provided in Methods.

### Initial testing and optimization of a monolithic agent in a stationary environment

As a reference point for subsequent evaluations, we first tested random and monolithic agents in a stationary environment, in which resource locations were fixed in the four corners of the grid (as in Figure 1c) (in all experiments, only one of each resource density was available, i.e. a single source of food). Maintaining homeostasis in this simple environment was not trivially accomplished, as evidenced by the performance of an agent that selected actions randomly on every time-step (Figure 2a). Despite chance encounters with resources, these were not sufficient to compensate for the depletion rates of the internal stats, which declined steadily over time. In contrast, a monolithic agent could be trained to maintain homeostasis for a variety of homeostatic set-points in this environment. Figure 2b summarizes the final internal stat level achieved after 30000 training steps (averaged over all 4 stats and over the final 1000 steps of training) for different set-points. The monolithic agent reliably achieved each set-point by the end of training. To ensure that subsequent comparisons were against the best possible monolithic agent, we selected a fixed set-point 5 for all stats (used in all subsequent experiments), and optimized the performance of the monolithic model by performing a search over performance-relevant hyper-parameters, such as the learning rate *α* and discount factor *γ*. Results of this hyper-parameter search (Figure S1) were used to select the best performing parameters for the monolithic agent. We then matched these parameters and others (including the total number of trainable parameters for each agent; see Table S1) between the monolithic and modular agents to compare them in the subsequent experiments.

**Fig. 2.**
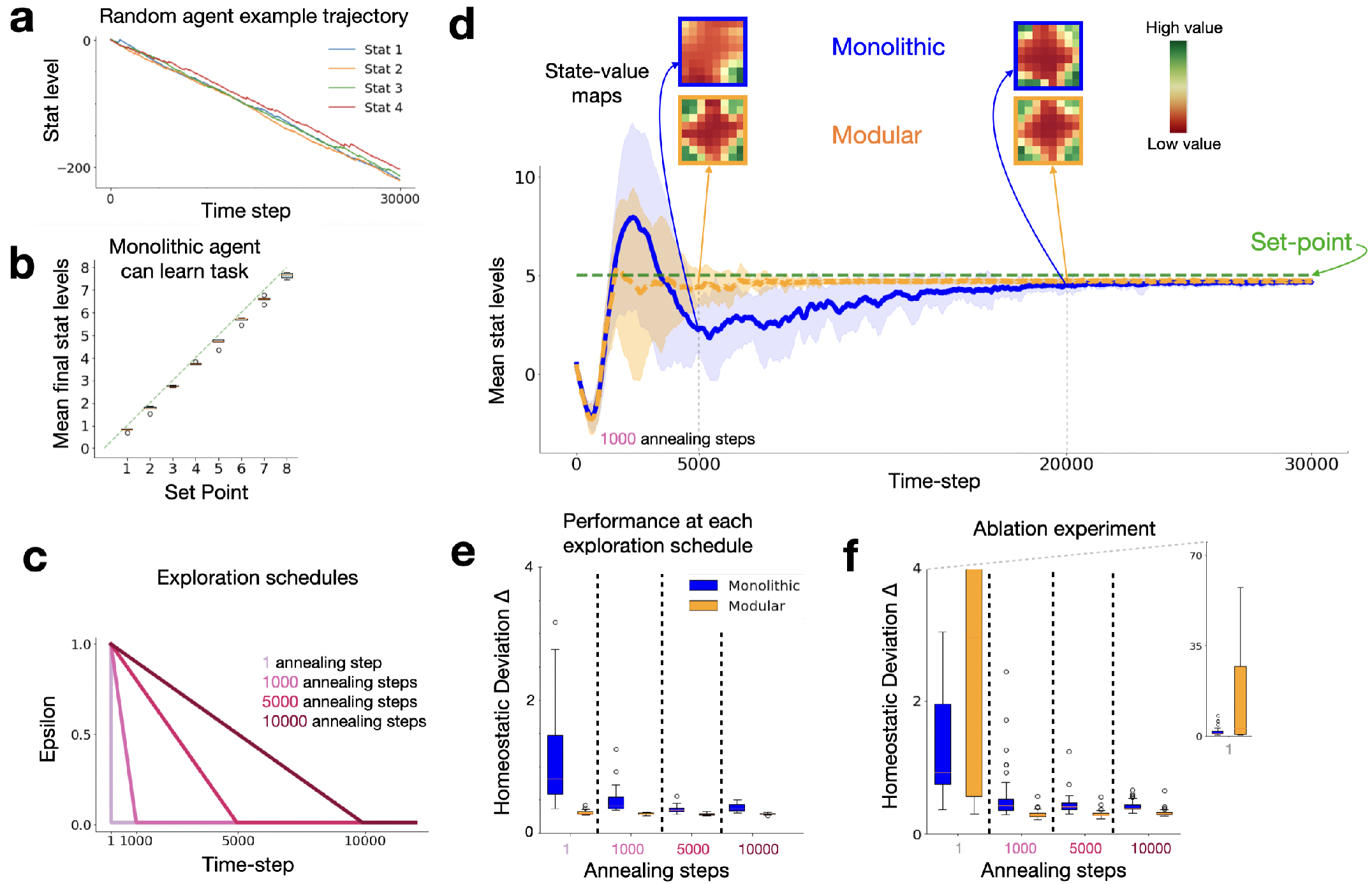
Performance of models in an environment with fixed resource locations. **a**: Stat levels decline over time for an agent that moved randomly on every time-step, shown for a period of 30,000 time-steps (same as period used for all subsequent experiments). **b**: Monolithic agents can learn to achieve homeostasis for a variety of set-points (data averaged for 10 agents; final stat levels calculated for each agent by averaging across levels of all four stats over the final 500 time-steps of training). **c**: Agents were further tested using 4 annealing schedules for *ϵ*-greedy exploration. Values of *ϵ* were initialized at 1 and linearly decreased on each time-step until reaching a final value of 0.01. This decrease occurred over a period of 1, 1000, 5000 or 10000 time-steps (i.e. schedule 1 applied effectively no annealing, while schedule 4 spread the annealing process over the first third of training). **d** Time course of average stat levels during training for monolithic (blue) and modular (orange) agents (*N* = 30 each; shading shows standard deviation at each time-point, and green dotted line represents set-point of 5 used for all four stats). Insets show representative heat-maps for the maximum action value in each state in the grid-world when all 4 stats were depleted equally to a value of 3 units for the for the monolithic and modular agents at time-points 5000 and 20000. The modular agent learned to represent the appropriate high value of all four corners of the grid (where the resources were located) much earlier in training. **e**: Direct comparison of the effects of exploration annealing on homeostatic performance of the monolithic and modular agents (*N* = 50 each) in an environment in which resources were fixed in the four corners of the grid-world, and decision noise was gradually decreased according to one of four annealing schedules. Homeostatic performance was calculated as in Methods Eq. 8 (lower is better). Performance of the monolithic agent (blue) relied heavily on the extrinsically imposed exploration, showing a substantial improvement in performance with increasing duration of annealing. The modular agent (orange) was essentially unaffected by the annealing schedule, achieving near maximal performance with virtually no extrinsically imposed exploration. Boxplots display inter-quartile range and outliers for *N* models. **f**: Results of an ablation experiment testing for the effects of intrinsic exploration in modular agents, in which action transitions were only saved into the memory of a particular module when that module took its preferred action, or when the action was selected randomly according to the epsilon schedule. As a control, for monolithic agents, saving transitions to memory was randomly skipped with a fixed probability (*p* = 0.3) to match the proportion of skipped experience in the modular agents. This rendered modular agents dependent on annealing, suggesting that intrinsic exploration was ‘knocked out’ and that performance was ‘rescued’ when extrinsic exploration was re-introduced.

### Modularity benefits exploration and learning

To investigate the impact of modularity on the need for exploration, we systematically varied the number of steps used to anneal *ϵ* (frequency of random action selection) in *ϵ*-greedy exploration from its initial to final value. We varied the *ϵ* annealing time for both types of agents over four schedules, in which exploration was reduced linearly from its initial value (*ϵ* = 1) to a final value (*ϵ* = 0.01) in 1 step or over 1K, 5K or 10K time steps (see Figure 2c). Figure 2d shows average stat levels over the course of training for the two types of agents in the 1K annealing schedule (See Figure S2 for example time-courses of separated stats). For both agents, stats briefly fell during the initial exploratory period, then stabilized toward the set-point. However, the modular agent consistently reached set-point faster, while the monolithic agent first overshot and then undershot the set-point before slowly converging on it, and exhibited substantially more variance while doing so (i.e., its stats reached more extreme values before converging). This overshoot-undershoot pattern has been previously observed (24), due to internal stats changing faster than the agent could adapt through learning. We investigated a range of other drive parameters, and found that the modular agent mitigated this pattern consistently (Figure S3).

The insets in Figure 2d show representative learned statevalue maps for each agent early (*t* = 5000) and late (*t* = 20000) in training. These maps were constructed, for testing at a given point in training, by externally and momentarily depleting all internal stats (setting them to a value of 3), and displaying the highest Q-value (for the modular agent, after summing action values across modules) in each grid location (high-valued states in green, and low-valued states in red), after which stats were returned to their previous value and training continued. In the very early stages of training (at 5000 steps), the modular agent learned a representation of values that reflected the state of the world (all four corners were valuable because all stats were depleted), whereas the monolithic agent had not. This pattern (i.e., modular agents learning appropriate representations very early in training) was observed in most training runs. In contrast, monolithic agents learned the appropriate value maps at much later stages in training (20000 steps).

Figure 2e summarizes the results of the different exploration annealing schedules, measured as mean deviation from set point (i.e., averaged over stats for each agent; see Methods). While monolithic agents gained progressively greater performance benefits from increasing periods of initial exploration, modular agents exhibited virtually no impact of extrinsically imposed exploration: With only 1 step of annealing, modular agents were already at or near maximal performance. This suggests that modular agents may have benefited from an intrinsic form of exploration, as a consequence of how action values were updated. These were updated for each sub-agent with respect to their individual objectives following every action, *irrespective* of which sub-agent contributed most strongly to the selected action. For sub-agents that did *not* contribute most strongly, the selected action could be considered (in mean expectation) to be approximately random (See Figure S4). This allowed those sub-agents to discover the values of actions that they would not otherwise have selected, providing an intrinsic and continual form of exploration.

To examine the potential contribution of intrinsic exploration to the performance of modular agents, we conducted an ablation experiment in which each module could only learn from actions that it *would* have taken if it had full control (that is, for which it was most responsible; see Methods for details). This manipulation substantially impaired the performance of modular agents in the absence of extrinsically imposed exploration, rendering them – like the monolithic agents – dependent on the annealing schedule, with performance ‘rescued’ as exploration was reintroduced with annealing (Figure 2f). In this ablation experiment, monolithic agents were also randomly deprived of learning from experience at a comparable rate as the modular agent as a control, and were relatively unaffected by the manipulation.

### The advantage of modularity is enhanced in changing environments and with more objectives

To increase task difficulty, we systematically varied two parameters: degree of non-stationarity (rate of change) in the environment, and number of homeostatic needs of the agent (and corresponding resources in the environment; see Figure S2 for example environments). We first introduced non-stationarity, such that each resource moved independently to a different randomly selected grid location at stochastic time intervals determined by a Poisson process with rate *λ*. Figure 3a shows results for monolithic and modular agents when *λ* = 0.02. The difference in performance between monolithic and modular agents was substantially greater in this environment compared to ones in which resources remained at fixed locations. Modular agents exhibited slightly greater reliance on exogenously applied exploration at the earliest points in learning; but this quickly diminished, and was dramatically less than monolithic agents which continued to exhibit heavy reliance on annealing.

**Fig. 3.**
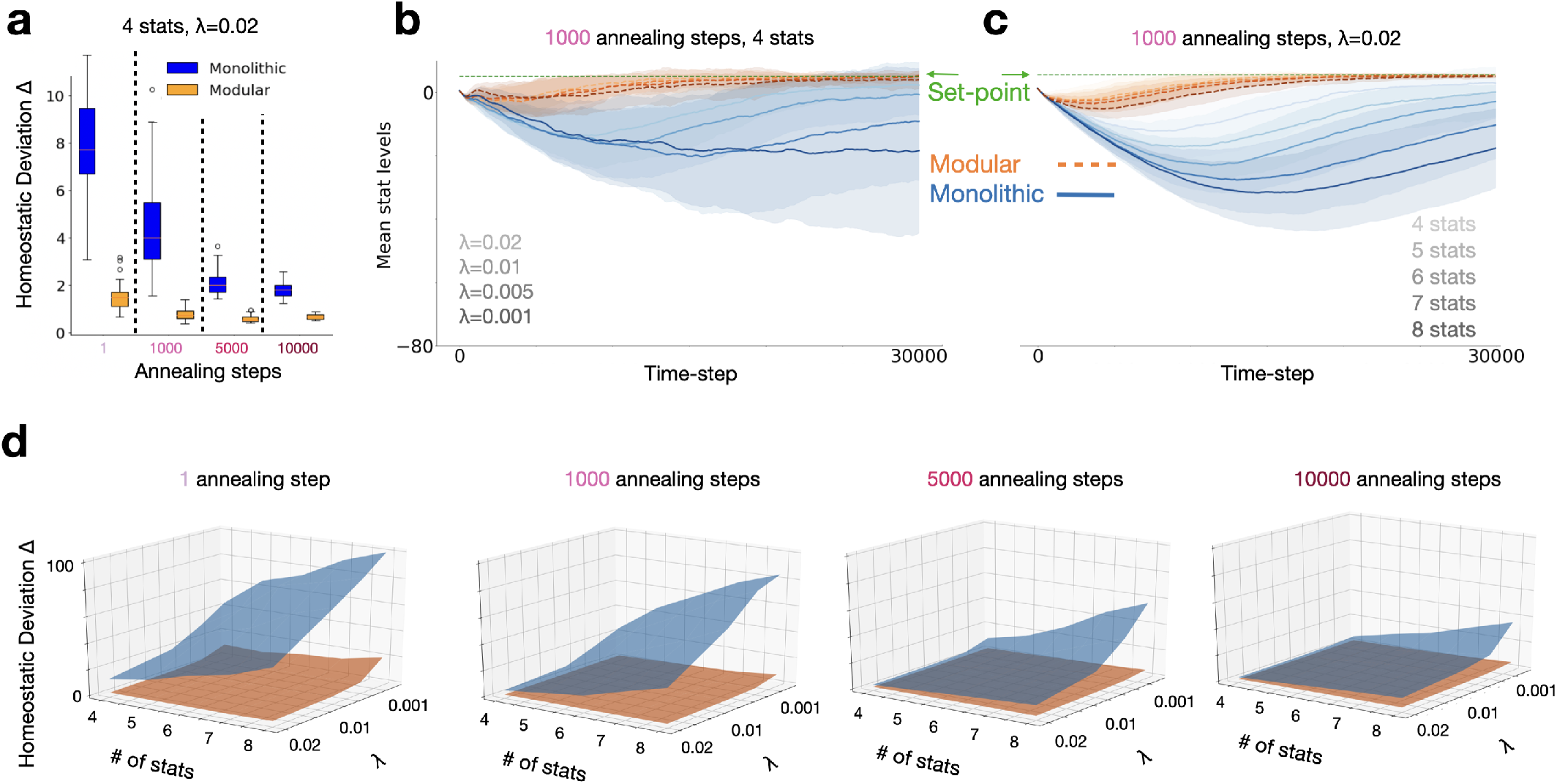
Performance of models in non-stationary and increasingly complex environments. **a**: Comparison of modular (orange) and monolithic (blue) agents (N=100 each) in an environment with four resources, each changing location at a Poisson rate of *λ* = 0.02. Homeostatic performance was calculated as in Methods Eq. [8] (lower is better). Modular agents were again relatively unaffected by the exploration annealing schedule, whereas the monolithic required significant exploration to have comparable performance (note that the differences between agents is greater here than in Figure 2e). Boxplots display inter-quartile range and outliers. **b**: Trajectories of stats levels (average of four stats) over training for modular and monolithic agents using the 1000 annealing step schedule, for environments with different rates of change varying from *λ*=0.001 (slowest, darkest line) to *λ*=0.02 (fastest, lightest line). For monolithic agents, homeostatic performance was worst in the most slowly changing environment, improving with rate of environmental change; whereas modular agents consistently and quickly achieved homeostasis regardless of the rate of change in the envinroment. **c**: Trajectories of stats levels (average of all stats) over training for modular and monolithic agents with different numbers of homeostatic objectives, from four (lightest line) to eight (darkest line), using the 1000 annealing step schedule in the fastest changing environment (*λ* = 0.02). For monolithic agents, homeostatic performance worsened with more homeostatic objectives, whereas the modular agent consistently and quickly achieved homeostasis regardless of the number of objectives. Shading represents standard deviation across all trained models.**d**: Summary of results across all manipulations. The duration of annealing (amount of extrinsic exploration) increase for plots from left to right, and the left and right horizontal axes of each plot represent number of stats and rate of resource location change (*λ*) respectively. The vertical axis displays homeostatic performance. The modular agent was able to robustly achieve homeostasis (orange surface remained mostly flat across all conditions).

We further tested the effects of non-stationarity by systematically varying *λ*. Figure 3b shows the time course for each type of agent (average of the four stats) over training (using the 1000 step annealing schedule) for different *λ*s, varying from the slowest rate of change (*λ* = 0.001, darkest lines) to fastest (*λ* = .02, lightest lines). Modular agents (orange) were largely impervious to the rate of change, consistently learning to achieve homeostasis relatively early during training. Monolithic agents (blue) performed worse overall, and performance worsened as resource locations changed more *slowly*. On further investigation, we found that agents occasionally fell into local minima solutions in non-stationary environments (possibly due to over-fitting on old resource locations), and that intrinsic exploration helped modular agents rely less on additional resource movements (which happened more often in fast changing environments) to escape these minima (See Figure S5 and related supplementary information). This suggests that, in slowly changing environments, the difference between modular and monolithic agents should be even greater when extrinsic exploration is decreased, an effect that is consistent with findings shown in the leftmost plot in Figure 3d (and discussed further below).

Finally, to compare how the two types of agents fared with an increasing number of objectives, we held *λ* constant at 0.02 and increased the number of stats and corresponding resources (and associated sub-agents in the modular architecture, while comparably increasing the number of trainable parameters in the monolithic model). Figure 3c shows the time course of training for 4 (lightest line) to 8 (darkest line) stats using the 1000 annealing step schedule. Again, modular agents achieved homeostasis regardless of number of objectives, even in this more difficult non-stationary environment, while the monolithic agent exhibited strong sensitivity to the number of objectives.

Figure 3d summarizes more results across all conditions. This highlights the observation that modular agents maintained good homeostatic performance (orange surface remains largely low and flat) across the three task manipulations studied: amount of extrinsically-imposed exploration (increasing from leftmost to rightmost plots), rate of non-stationarity (right axis of each plot), and number of homeostatic needs (left axis of each plot); whereas the performance of monolithic agents was sensitive to all of these, with performance degrading in a parametric fashion in response to each manipulation (blue surface overall higher and steeply pitched). A more complete set of time-courses is provided in Figures S6 and S7, and we support the generality of this effect by replicating it with a varied exploration procedure (Figure S10) and over a range of different drive parameters (Figure S3). Note that the difference between modular and monolithic agents in slowly changing environments (*λ* = 0.001) was greatest with the least amount of extrinsic exploration (leftmost plot) and decreased with longer annealing, whereas the modular agents showed good performance and little change as a function of extrinsic exploration, consistent with the suggestion above that intrinsic exploration helped them escape local minima in slowly changing environments.

### Internal representations learned by monolithic and modular agents

In addition to differences in dependence on exploration, another important difference is the types of value representations each agent is capable of learning. Figure 4a provides a conceptual illustration of the point that, for the monolithic agent, there will be a loss of information when separate reward components are combined into a single scalar value (the flat black line is the sum of the four sources of reward as they vary over space or time), which can make it difficult to learn a policy sensitive to the individual objectives. In contrast, in the case of the modular agent, policy learning for each objective was kept separate and based only on the reward for the corresponding objective. The consequences of this difference are shown in Figure 4b. In this example, a monolithic and a modular agent were each trained in the same environment with non-stationary resources, and then their state values and policies were examined immediately after changing all of the resource locations, by setting all stats to their set point except for one, corresponding to the resource now placed at the center, that was set to 0 (i.e., signifying the greatest need). The color coding of the state-value maps (at the far right of the figure) shows that both agents accurately identified the location of the resource with the greatest current value (green at the center of the maps). However, the policies learned by the agents (shown as arrows in each grid location indicating the preferred action at that location) differed considerably between them. The modular agent learned a policy by which, starting at any location, a path following the policy lead to the resource with the greatest current value (at the center). In contrast, for the policy learned by the monolithic agent, many paths failed to lead to that location. This difference was because the monolithic agent had the more difficult task of constructing a global policy for every possible set of needs and resource locations (that is, a form of conjunctive coding that grows exponentially with the number of resources), whereas the modular agent constructed individual policies for each objective (a form of compositional coding, that grows only linearly with the number of resources), that could then simply be summed together (shown in the lower middle plots) to allow the policy associated with the most valued resource to have the greatest influence.

**Fig. 4.**
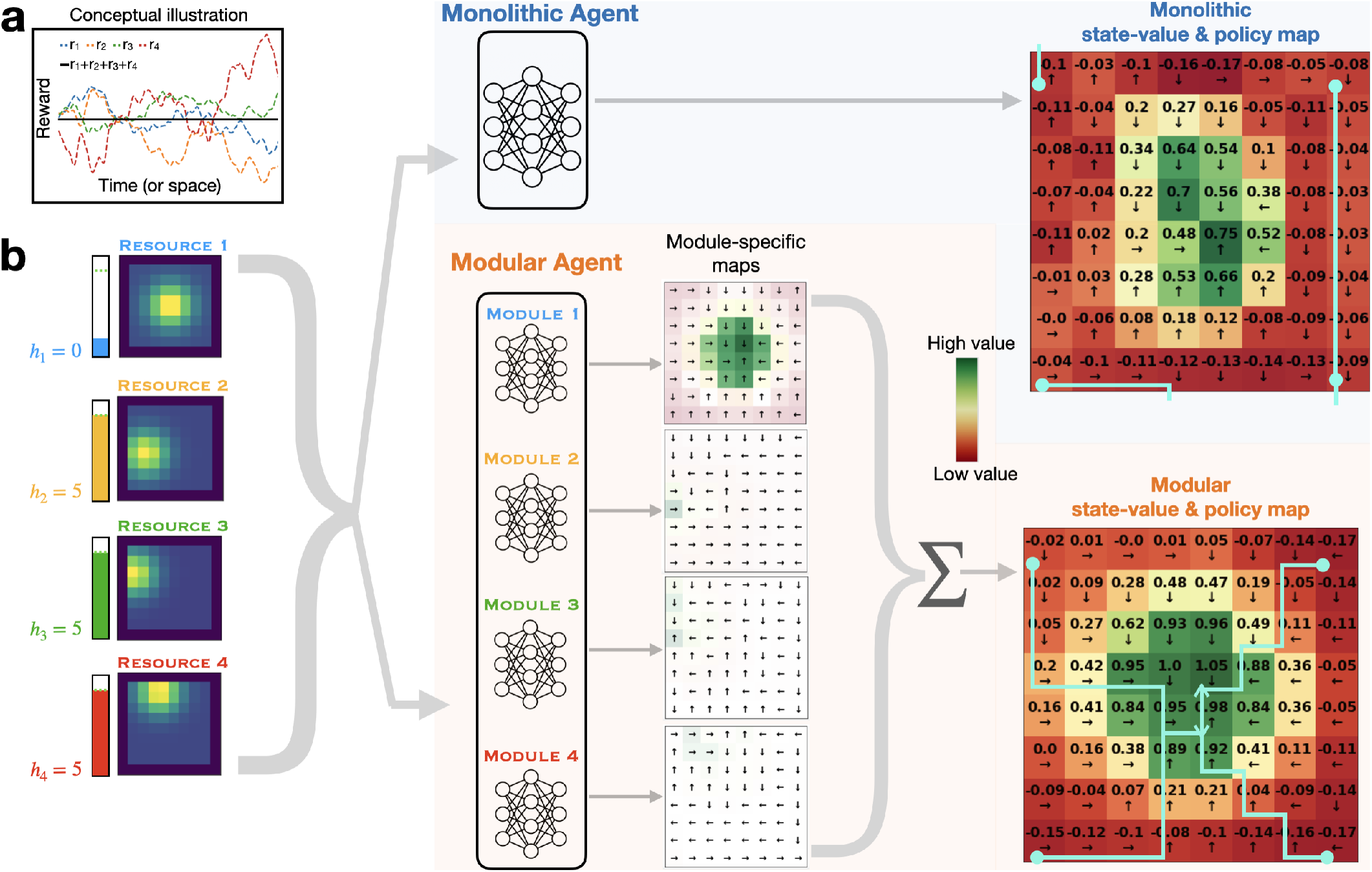
State-values and policies learned by monolithic and modular agents. **a**: Loss of information in monolithic architecture. Plots of hypothetical time course for rewards relevant to each of four sub-agents, that sum to a constant value at every time point. This extreme case highlights the potential loss of information available to a monolithic agent: summing the reward signals obscures differences in the reward relevant to each objective, making it difficult for it to develop a policy that is sensitive to them individually. **b** State-value and policy maps learned by each type of agent. A monolithic and a modular agent were trained for 30000 steps in the same non-stationary environment (*λ* = 0.02). After training, all resource locations were changed, with resource 1 moved to the center of the grid (maps at left), and the internal stat for that resource *h*_1_ depleted to 0, while all other stats were set to their set-point of 5 (level meters to the left of the resource maps). This state was then passed to both models, to construct state value and policy maps (monolithic in upper right; modular in lower right), which display the maximum Q-value output for each grid location and its corresponding action direction (black arrows). For the modular agent, maps in the middle show the values outputted by each individual module (with color reflecting *relative* state values within each map, and transparency the *absolute* value at each grid location allowing comparison of values between modules). Module 1 had higher action values overall, reflecting the depleted state of stat 1, and thus contributing the most to the final value map (at right) after summing action values across modules. The final value/policy maps show that the modular agent had a clear path to resource 1 from any location in the grid. However, the monolithic agent, while recognizing the higher value of resource 1, had many paths that did not lead there. Paths can be traced by following the black arrows (some example paths starting from grid corners are shown in light blue).

## Discussion

In this article, we showed that a modular architecture can learn more efficiently and effectively to simultaneously manage multiple homeostatic objectives, both in non-stationary environments, and as the number of independent objectives was increased. We provided evidence that this was for at least two reasons: intrinsic exploration and a factoring of representational learning by objective. Intrinsic exploration reflected an emergent property of the modular architecture, in which the actions determined by the needs of one sub-agent served as a source of exploration for the others, allowing them to discover the value of actions they may not have otherwise chosen in a given state. Modularity also allowed each sub-agent to focus on, and “specialize in serving” its own objective, allowing it to learn objective-specific policies that could be invoked as a function of their current value in determining the action of the agent as a whole. Together, these factors contributed to the ability of the modular architecture to support rapid adaptation in changing environments (even when change was infrequent), and to deal gracefully with the “curse of dimensionality” associated with an increase in the number of objectives. In the remainder of the article, we consider how our findings relate to relevant ones in reinforcement learning, biology, neuroscience and psychology.

### Connections to reinforcement learning

#### Modularity

Some have argued that reward maximization in RL is sufficient to produce all known features of intelligent behavior (41). In practice, such a reward function, even if possible to specify, is difficult to maximize, because it requires gathering enough information to map all states of the world onto a preferred action. The alternative that we have considered here – learning separate, more tractable sub-problems in parallel – is not a new one. Both earlier (30, 42–47) and more recent (31, 33**?** –37) work has studied ‘modular RL,’ in which modules are specialized for particular objectives in the tradition of mixture-of-experts systems (48). In most of this work, and that presented here, modules compete for action in parallel. Other kinds of modular organization exist, such as the option framework in hierarchical reinforcement learning (HRL (49)), in which action modules are arranged in a strict temporal hierarchy. However, in HRL modules are still all under centralized control. From this perspective, our organization is heterarchical or decentralized. These approaches may be complementary. For example, one could conceive of objective-specific modules being separate HRL agents in their own right, or alternatively, that certain objectives might occasionally assume a hierarchical role over others. The world contains structures that are both tree-like (i.e., hierarchical) as well as independently varying (heterarchical) (50) and, accordingly, effective learning agents are likely to reflect both kinds of organization.

#### Multiple objectives

Our work is also related to the field of multiobjective reinforcement learning (MORL), which has studied how to optimally learn multiple policies that cover a range of trade-offs between objectives, known as a Pareto front (13, 39, 51). Typically, MORL learns policies that do not focus on single objectives, but rather specific preferences over them (i.e. it would learn a separate policy for valuing food and water equally, for valuing food twice as much as water, etc.). However, since all objectives must be considered together, it is subject to problems similar to those inherent to the monolithic approach – the dimensionality of the problem grows with the number of objectives – and to the challenges of non-stationarity, as the relative importance of objectives is likely to change over time. As we have shown here, the modular architecture has the potential to deal gracefully with these problems. The field of multi-agent RL has also studied multiple objectives, but typically in the setting of separate entities competing or cooperating in a shared environment (52); in contrast, we have considered the benefits of multiple sub-agents competing within a single agent (i.e., body), in which they compete for control of action rather than, or in addition to resources.

#### Non-stationarity

Non-stationarity is a fundamental feature of the world, and RL usually requires specialized machinery to deal with it, such as adaptive exploration based on detecting context changes (28, 53–56). In contrast, the modular architecture benefits from emergent exploration that arises from the competition between sub-agents. Additionally, what counts as a change in the environment might differ between modules. We suggest that equipping modules with learnable attention masks (see Methods) allowed them to ignore changes in the environment that were irrelevant to their particular goal; this raises the interesting possibility that empirical learning phenomena, such as latent inhibition (57) and/or goal-conditioned attention (58), may reflect a similar form of learned inattention in the service coping with non-stationarity.

Recent work has also suggested decomposing a reward signal into separate components as a strategy to deal with non-stationarity in the reward function itself (e.g., to account for the fact that water becomes more rewarding when thirsty). In this ‘reward bases’ model (59), the importance of each learned component of value can be re-weighted on the fly, allowing for immediate adaptation to the changing physiologic state of the agent. In the modular architecture we considered here, agents directly sensed their own physiologic state. While this rendered the reward function itself stationary (i.e. the reward associated with any specific combination of the state of the agent and the environment was constant), we consider this a more plausible design, in which the capacity to dynamically re-weight module importance as a function of agent state and/or change in the environment was implicit and learned, as opposed to being externally imposed.

#### Exploration

As already noted, the need for exploration is a fundamental feature of – and challenge for – RL, and existing solutions generally make use of some form of noise or explicit exploratory drives/bonuses (27). The work presented here suggests an additional class of strategy: that exploration can arise as an added benefit of having multiple independent objectives, since exploitation from the perspective of one is exploration from the perspective of others. This form of exploration is analogous to that described by (60), in which hierarchical organization of policies provides ‘semantic exploration’ that arises from actions at different time-scales. In this article, heterarchical organization of policies provided semantic exploration arising from actions serving different objectives. A similar architecture is also described by (61), in which multiple DQN modules with the *same* objective provided exploratory noise (i.e. from different random initialization of module networks). In the architecture presented here, we have suggested that exploration was driven primarily by diverse objectives rather than diverse initializations. Finally, while explicit exploration could be combined with a modular architecture by implementing a dedicated “exploration module” (62), an appealing feature of the modular architecture on its own is that exploration arises as emergent property of the system. That said, this does not preclude the possibility that natural agents make use of both strategies (63). Relatedly, exploration may also emerge naturally from the free energy principle, in order to reduce model uncertainty (64), or as a result of Bayesian model averaging, in which actions are derived from a weighted sum over policies that may vary in complexity or reliability (65). Our findings reflect a mechanism in modelfree systems, in which uncertainty is reduced by virtue of policy diversity along the dimension of organismal need (rather than model complexity), but which could be complemented by the more complex world-models learnable in the free energy framework and/or model-based RL.

### Connections to biology and evolution

In order to survive, organisms must maintain a set of physiological variables within a habitable range, with the capacity of these variables to vary independently introducing the potential for conflict between them. Frameworks to explain homeostasis include drive reduction theory (66), predictive control (67), active inference (68), motivational mapping (69) and more recently homeostatically regulated reinforcement learning (HRRL) (24). HRRL is a general framework that aims to account for the others, and has been used not only to explain low-level drive satisfaction but also more abstract goal-seeking (25) and even psychiatric phenomena (70). However, HRRL collapses a high-dimensional reward surface into a single number, and its challenges as a monolithic approach have not been previously interrogated through simulation. Our work suggests that maintaining an independent set of homeostatic reward components offers computational advantages that are in keeping with modularity as a general principle in biological organization.

Organisms exhibit modularity from the level of organelles and cells to organ systems and brain regions. While modularity exists on a continuum and its definition is non-trivial, especially in learning systems (71), having components that function independently to some extent has been shown to be favoured by evolutionary processes. For example, neural networks evolved with genetic algorithms develop modularity when trained on goals that change in an independent fashion, whereas monolithic networks develop for stationary goals (72). Furthermore, when neural networks are evolved simply to minimize connection costs, they both become increasingly modular and adapt better to new environments (73). In bacteria, metabolic networks are more modular the more variable are their environments (74). These findings are consistent with our results that modularity provides significant performance benefits in non-stationary environments. More broadly, we might even expect to find similar adaptations at the population level, especially in complex societies of organisms; this may explain why we see humans with widely varying or competitive attitudes within groups, even when more homogeneous cooperative populations might be expected to result from group selection (75).

### Connections to neuroscience

There is growing evidence for modular RL in both brain and behavior. It has long been recognized that the most basic homeostatic needs (e.g., osmotic balance, metabolic state, thermoregulation, etc.) are represented in compartmentalized form within the hypothalamus (76), and similar modular organization has been reported within higher level structures. For example, within the basal ganglia, striosomes have been proposed as specialized units that compete for action selection (77), which is consistent with recent evidence for heterogeneous dopamine signaling and mixture-of-experts learning in that region (78). Diverse dopamine responses coding for salience, threat, movement, accuracy and other sensory variables (79) have also been observed, including sub-populations of dopamine neurons that track distinct needs related to food and water (80) and social reward components (81). Behavioral modeling has suggested modular reward learning for separate nutritional components like fat and sugar (82). While it is possible that such heterogeneity is simply a side effect of standard scalar reward maximization given uneven sensory inputs to dopamine neuron populations (83), our work suggests the possibility of a more functional role, predicting that dopamine heterogeneity may track the multiplicity of ongoing needs an agent must satisfy, and distinct learning systems associated with these.

There is also mounting evidence for separated value functions in the human brain (84, 85), and that decision dynamics are best modelled using competing value components (86). Others have proposed that opposed serotonin and dopamine learning systems reflect competition between optimistic and pessimistic behavioral policies (87), and that human behavior can be fit best by assuming it reflects modular reinforcement learning (88, 89). Our work provides an explanation for these findings, which are all consistent with the idea that different objectives compete for behavioral expression in parallel (90). There is also evidence for hierarchically-structured value signals in the brain (91, 92). That said, and as noted above, where human brain function should be placed along the continuum of centralized to distributed control of behavior remains an interesting and important open question (93).

### Connections to psychology

The study of intrapsychic conflict has a long history in psychology; between ego, id and superego (14), opposing beliefs (94), approach and avoid systems (95), affects (96), motivations (97), and even sub-personalities (98). Despite their specifics, these accounts all have in common the following: the assumption that an individual is comprised of multiple distinct subsystems responsible for satisfying different objectives (i.e., “multiple selves”), all of which compete to express their proposed actions in the behavior of the individual (whether these be covert “actions,” such as internal thoughts and feelings, or overt physical actions). While there have been a large number of theories that propose a qualitative account of the mechanisms involved, to date there have been few formally rigorous or quantitative accounts, nor any that provide a normative explanation for why the mind should be constructed in this way. In cognitive psychology, theories and mechanistic models have been proposed regarding the conflict between controlled and automatic processing (99, 100) and, similarly in cognitive neuroscience, between model-based and model-free RL systems (101) as well as the role of conflict in processing more generally (102). However, none of these deal explicitly with conflicting policies that have altogether different objectives (i.e., the goal has typically been to maximize a single scalar reward), nor have any provided an account of why conflict *itself* might actually be intrinsically useful (i.e., for exploration). Our work provides a normative account of these factors.

In the psychodynamic literature more specifically, not only has there been an effort to describe conflict between opposing psychological processes, but also its resolution by “defense mechanisms” (6, 103–106), which we speculate might reflect complex ways to arbitrate between modules. In addition, a psychotherapy often aims to “integrate” psychological sources of conflict (107–109). Such integration might have to do with a developmental process that progresses along the previously mentioned continuum from heterarchical to hierarchical organization. Our work is certainly far removed from concepts like defense mechanisms or psychological integration, but we hypothesize that modular RL is a framework upon which more formal correspondences to the conflict-based psychodynamic theories could eventually be built.

### Limitations and future work

There are several limitations of the work presented here, that could be addressed in future research. First, the reward decomposition across modules was pre-specified in a simple way: one sub-agent for each resource. However, in nature, correlations can exist between internal stats (i.e. food replenishing nutrients and water). Early work suggests that related drives (110) interact behaviorally more than less related drives (111), hinting that policy modularity may vary with drive independence. Future work would therefore relax the one-module-per-drive constraint, and explore performance in environments with different correlation structures. For example, given three correlated and two uncorrelated drives, a hybrid model with one 3D reward component and two 1D reward components might out-perform a fully monolithic or fully modular agent. To what extent such organization is innate or learned is another open question.

Second, our task definition did not capture the true multiplicity of objectives humans face, and their variety (for example, one could imagine a pain module, contributing only negative rewards, or modules for more abstract objectives, such as for money or social respect). While our results concerning the curse of dimensionality were encouraging in considering an agent with a large or expandable set of objectives, we modelled all objectives identically with reward functions that were symmetric around a set-point. Furthermore, although rewards derived from drive reduction may not be sufficient to capture all objectives, we predict that the benefits of modularity would extend to different reward functions to the extent that sources of reward are independent, as previously discussed.

Next, we did not explore model-based approaches in this work. Such approaches may have significant promise; a modelbased but unidimensional version of HRRL was able to account for features of addiction (112), and a single-agent model-based approach demonstrated impressive capabilities in balancing multiple homeostatic goals in changing environments (113). Nevertheless, such model-based approaches may benefit from modularity in order to avoid interference between tasks (114), and to deal with curse of dimensionality in complex environments. Future work should therefore explore hybrid systems in the rich space that combines model-free, model-based, monolithic and modular learning components.

Finally, a critical feature of any modular architecture is the mechanism used for adjudication. We used a minimalist implementation that simply summed Q-values, as a way of identifying the benefits intrsinsic to the modular structure, rather than the adjudication mechanism. However, more sophisticated forms of arbitration, that could flexibly and dynamically re-weight individual module outputs, might better exploit the benefits of modularity. This would be important for cases in which different objectives may change in relative importance faster than modules can adapt through reinforcement learning, or if more global coordination is required for certain tasks.

## Concluding remarks

The question with which we began was: How do agents learn to balance conflicting needs in a complex changing world? The work presented in this article suggests that a modular architecture may be an important factor, addressing two critical challenges posed by the question: the ability to adapt effectively and efficiently as needs and the resources required to satisfy them change over time; and the ability to avert the “curse of dimensionality” associated with an increasing number of objectives. These observations may help provide insight into the principles of human brain organization and psychological function, and at the same time inform the design of artificial agents that are likely to a face similar need to satisfy a growing number of objectives.

## Materials and Methods

### Approach

We draw on the field of reinforcement learning, which models learning agents in a Markov decision process (MDP) in the following way: At each time-step *t*, an agent perceives the state of the environment s_*t*_, takes an action *a*_*t*_ based on its learned behavior policy *π*(*s*), causing a transition to state *s*_*t*+1_ and receiving reward *r*_*t*_. The agent tries to collect as much reward as it can in finite time. However, since its lifespan is both unknown to the agent and long relative to the timescale of learning, the time horizon for reward collection is typically treated as infinitely far away. The agent thus aims to maximizes the sum of discounted future rewards, defined as the return *G*_*t*_ [1],

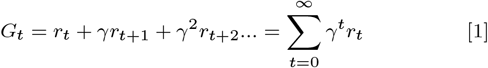

where γ ∈ {0, 1} is the discount factor, a parameter that determines the present value of future rewards. The agent can then learn to estimate the expected return of action *a* in state *s*, defined as the action value *Q*(*s, a*) (equation [2]), which we sometimes refer to simply as *Q*-value or action value for clarity.

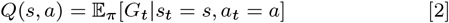

Once the agent has learned to estimate action values accurately, then a good behavioral policy simply takes the action with the highest Q-value, also called the greedy action, in each state [3]:

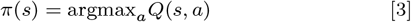

However, considering that Q-values are sub-optimal while learning takes place, always choosing actions greedily ensures that potentially more rewarding actions are never tried out. The simplest way to approach this explore-exploit trade-off is to select actions greedily with probability ϵ but randomly otherwise (termed ϵ-greedy exploration). Then, using the agent’s experience, we use the Q-learning algorithm (115) to update action values in order to minimize the magnitude of the temporal difference (TD) error δ defined in [4].

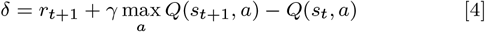

In the case where there are a large, or infinite, number of states (i.e. because state variables are continuous), then function approximation is required to learn a mapping Φ(*s*) from states to action values. Typically a neural network is used, with parameters *θ* that can then be learned by gradient descent to directly minimize the TD loss function. Such networks are termed deep Q-networks (DQN) (40), and standard implementation details are summarized in Training Details below.

Lastly, we summarize how we apply RL in the context of multiple goals. Given a set of objectives {*o*_1_, *o*_2_, …*o*_*N*_}, we define a vector of rewards 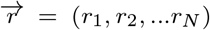 corresponding to each objective, and distinguish our two main approaches in general terms. The monolithic approach computes a scalar reward 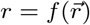 and then learns Q_monolithic_ as usual, whereas the modular approach first learns a corresponding vector of Q-values 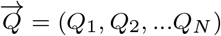 and then computes 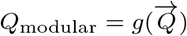, for some functions *f* and *g*.

While many other reinforcement learning algorithms exist, such as those that learn models of the environment (116), or those that dispense with Q-values entirely (117), we focus our work on model-free Q-learning because of its simplicity, interpretability, empirical support from neuroscience (118) and success as an off-policy algorithm. Off-policy learning means that agents can learn from experience that did not derive from its current policy, and is important because in this work, modules learn in parallel from a shared set of actions.

### Environment

We developed a flexible environment in which an agent could move around and collect resources in order to fulfill a set of internal physiologic needs. Specifically, we constructed an 8x8 grid-world, where each location (*x, y*) in the grid indexed a vector of resource abundances of length *N* (i.e., there were *N* overlaid resource maps). For example, in an environment with 4 resources, the grid location (1, 1) might have resource abundances of [0, 0, 0.8, 0], indicating 0.8 units of resource 3 was present at that location. The spatial distribution of the amount of each resource was specified by a 2D Gaussian with mean *µ*_*x*_, *µ*_*y*_ and co-variance matrix Σ, and was normalized to ensure there was always the same total amount of each resource (see Figure 1a).

The agent received as perceptual input a 3x3 egocentric slice of the *N* resource maps (i.e. it could see the abundance of all resources at each position in its local vicinity, as well as a wall of pixels set to -1 if it was next to the border of the grid). In addition to the resource landscape, the agent also perceived a vector *H*_*t*_ = (*h*_1,*t*_, *h*_2,*t*_, …, *h*_*N,t*_) consisting of *N* internal variables, each representing the agent’s homeostatic need with respect to a corresponding resource. We refer to these variables as “internal stats” or just “stats” (such as osmostat, glucostat, etc.) which we assume are independent (*h*_*i*_ is only affected by acquisition of resource i) and have some desired set-point 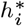. One should imagine these variables in the most general sense - while our study used a homeostatic interpretation, one could imagine the internal variables also representing more abstract notions like distance to a particular goal (70). For most of our experiments, all homeostatic set-points were fixed at the same value 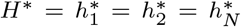 and did not change over the course of training.

To interact with the environment, on each time-step, agents could move to a new location by selecting one of four cardinal directions on the grid. For each internal stat, if the abundance of resource *i* in the new location was above some threshold *R*_thresh_, the stat *h*_*i*_ increased by that abundance level. Additionally, each internal stat decayed at a constant rate to represent the natural depletion of internal resources over time (note: resources in the environment themselves did not deplete). Thus, if the agent discovered a location with a high level of resource for a single depleted stat, staying at that location would optimize that stat toward its set-point, however others would progressively deplete. Agents were initialized in the center of the grid, with internal stats initialized below their set-points at a value of 0, and were trained for a total of 30, 000 steps in the environment. Thus, the task was for agents to learn in real time to achieve homeostasis, in a continuous-learning infinite-horizon setup (i.e. the environment was never reset during training, to more closely reflect the task of homeostasis in real-world organisms). Environmental parameters are summarized in Table S1.

### Models

In order to succeed in the environment just described, any RL agent would need to learn a function that converts its perceptual input into a set of optimal action values. Strategies to do so range from filling in a look-up table that maps all possible states to their values, to training a neural network to approximate such a mapping function. Our set-up required the latter strategy, pragmatically because homeostatic variables were continuous and therefore could not be tabulated, and theoretically because we believe the brain likely uses this kind of function approximation for reward learning (119).

### Monolithic agent

We created a monolithic agent based on the deep Q-network (DQN) (40). The agent’s perceptual input at each time step was a concatenation of all local resource levels in its egocentric view, along with all internal stat levels. Its output was 4 action logits subsequently used for ϵ-greedy action selection. We used the HRRL reward function (24) which defined reward at each time-step *r*_*t*_ as drive reduction, where drive *D* was a convex function of set-point deviations, with convexity determined by free parameters m and n; see equation [5].

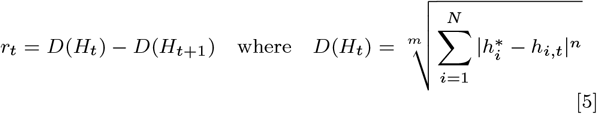

### Modular agent

We created a modular agent that consisted of a separate DQN for each of *N* resources/stats. Here, each module had the same input as the monolithic model (i.e. the full egocentric view and all *N* stat levels), but received a separate reward r_*i,t*_ derived from only a single stat. The reward function for the ith module was therefore defined as in equation [6], where drive *D* depended on the *i*th resource only. This 1-dimensional version of HRRL has been used previously (112), so that maximizing the sum of discounted rewards is equivalent to minimizing the sum of discounted homeostatic deviations for each module individually.

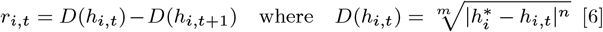

To select a single action from the suggestions of the multiple modules, we used a simple additive heuristic, based on an early technique called greatest-mass Q-learning (43). We first summed Q-values for each action across modules, and then performed standard ϵ-greedy action selection on the result. More specifically, if *Q*_*i*_(*s, a*) was the Q-value of action a suggested by module *i*, a greedy action *a*_*t*_ was selected as in equation [7].

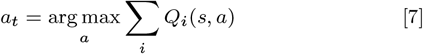

### Drive parameters

We made a principled selection of the drive parameters that would be suitable for both agents. First, the constraint *n* > *m* > 1 was necessary to be consistent with physiology (24); for example, drinking the same volume of water should be more rewarding in states of extreme thirst compared to the satiated state. Second, we wanted both agents to have a similar drive surface topology. Therefore, we selected (*n, m*) = (4, 2), such that 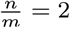, providing the modular agent with a quadratic drive surface.^*m*^Other parameter settings were investigated in Figure S3.

### Training details

All Q-networks were multi-layered perceptrons (MLP) with rectified linear nonlinearities, trained using double Q-learning (120) (a variant on standard deep Q-learning that uses a temporal difference Huber loss function, experience replay and target networks (40)). More specifically, on each time-step of training, a transition consisting of the current state, action, next state, and reward were saved into a memory buffer. On each time-step after a minimum of 128 transitions had been saved, a batch of at least 128 and up to 512 transitions was randomly sampled from memory, and used to backpropogate the TD-loss from equation [4] through the network. Gradient updates were performed on network parameters using the Adam optimizer (121). Importantly, we matched the number of trainable parameters between the modular and monolithic agents (to ensure that if the modular agent consisted of *N* DQN modules, it did not have *N* times as many parameters as the monolithic model, possibly giving it an unfair advantage).

For both models, ϵ was annealed linearly from its initial to final value at the beginning of training at a rate that was experimentally manipulated as described in the main text. To quantify model performance, we calculated the average homeostatic deviation per step ∆ between two time points *t*_1_ and *t*_2_ after a sufficient period of learning and exploration as in equation [8] (Lower ∆ therefore indicated better homeostatic performance). We used *t*_1_ = 15k and *t*_2_ = 30k, representing performance across the second half of the training period (except for results presented in Figure 3a which used *t*_1_ = 25k for visualization purposes).

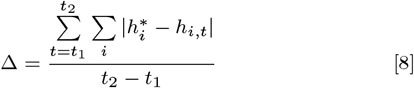

Finally, in non-stationary environments, learning a sparse attentional mask over the input to each network was required for good performance (see Figure S12). Therefore, all models in all environments were trained with such a mask. For example, assuming the input *I* to our networks was a vector of length 40 (i.e. consisting of four 3x3 egocentric views and four internal stats), and *A* was a 40 element masking vector, then the input was element-wise multiplied by the masking vector before being passed to the network as *A* ⨀ *I*. All elements of *A* were initialized to a value of 0.1, and were optimized along with the rest of the network during training (for the modular agent, each module learned a separate mask). An L1 regularization term applied to *A* was added to our loss function so that loss *L* = *δ* + *β* Σ_*i*_ |*A*|, where *A*_i_ were individual elements of *A, β* was a weighting parameter, and *δ* was the TD-loss already described. Additional parameters for both models as well as the environment are summarized in Table S1.

### Ablation experiment

To investigate the sources of exploration in the modular agent, we performed an ablation where experienced transitions were only saved to memories for a particular module in the following cases: a) the action taken was non-greedy (i.e. random) or b) the action taken was the preferred action of that module. In the monolithic case, in order control for less transitions being stored overall, 30% of non-greedy actions were randomly selected to not be stored in memory, an amount that was roughly similar to the number of transitions that were not saved for the modular model.

## ACKNOWLEDGMENTS

This project / publication was made possible through the support of a grant from the John Templeton Foundation. The opinions expressed in this publication are those of the authors and do not necessarily reflect the views of the John Templeton Foundation. This work was also supported in part by the Office of Naval Research.

## Supplemental Information

### Hyperparameter optimization

We performed our initial hyper-parameter search to optimize the monolithic model in an environment with 4 resources that were fixed in the 4 corners of our grid-world. We first adjusted hyperparameters by hand, in order to learn that the learning rate α and the discount factor γ were the most important factors when it came to overall performance. We then performed a grid-search over values α ∈ [1*e* − 5, 5*e* − 5, 1*e* − 4, 5*e* − 4, 1*e* − 3, 5*e* − 1] and γ ∈ [0, 0.1, 0.2, 0.3, 0.4, 0.5, 0.6, 0.7, 0.8, 0.9, 0.99]. We found that the best setting was α = 1e − 3, γ = 0.5 as in Figure S1. These values were then used for all models and experiments. Other hyper-parameters were adjusted by hand but stayed near typical values, and can be seen in Table S1 for the case of 4 resources. In order to maintain matched number of parameters for greater number of resources / modules, we added the following number of hidden layer units to the baseline 1024 units of the monolithic model: 0,130,250,360,470 for 4,5,6,7,8 modules respectively.

### Example trajectories

We visualize randomly selected example trajectories in Figure S2 for three different environmental set-ups -4 stationary resources, 4 non-stationary resources, and 8 non-stationary resources. All environments are non-trivial; agents moving randomly deplete their stats monotonically. Modular agent examples display how all stats eventually reach set-point in all environment set-ups. Monolithic agent examples struggle to varying degrees.

### Drive Parameters

Different drive surface parameters represent different reward function properties. For example, in the monolithic case, the greater the coefficients (*n, m*), the larger the relative size of multi-dimensional drive reductions relative to unidimensional ones, which become equal when n = m, and reverse in importance when (*n, m*) < 1. As well, with *n* > *m* > 1, the drive gradient is greater farther from set-point. With n < m < 1, the drive gradient is greater closer to set-point. Therefore, we investigated an expanded set of drive parameters which increased the generalizability of our results while offering additional insights into how the shape of the reward function affected the behavior of our agents. We tested parameters (n,m) = (4,2), (3,2), (2,2), (1,1), and (0.8,0.9). (Note that (2,2) and (1,1) are degenerate with respect to the modular model, but we included both anyway).

Figure S3 (left) replicates Figure 2d (stationary environment) for the new parameter settings, with corresponding drive surfaces for reference (2D for monolithic and 1D for modular). The modular agent maintained a very similar trajectory in all settings (achieved homeostasis early in training and remained near set-point with low variance). The overshoot-undershoot pattern in the monolithic agent was preserved, but diminished as (*n, m*) decreased in a way that worsened performance (i.e. for (*n, m*) = (0.8, 0.9), the monolithic stats had higher variance and had trouble reaching set-point). Therefore, when the drive surface did not favour multi-dimensional drive reductions, or even favored unidimensional reductions, the monolithic agent did not become equivalent to the modular one.

As well, when the drive gradient was steeper near set-point (*m* = 0.8, *n* = 0.9), it appears that caused instability (since both positive and negative rewards would be higher), and less drive towards set-point when farther away from it, causing higher stat variance and worse homeostatic performance overall in the monolithic agent. Speculating for the modular case, we wonder whether ‘negative feedback’ from other modules helped to both ‘dampen’ the overshoot-undershoot, while also preventing the instabilities near set-point that we see in the monolithic case for lower parameter values. Lastly, Figure S3 (right) generalizes our results from Figure 3d (non-stationary environment) over these new drive parameters for two extremes of exploration schedules. Again, the modular agent exhibits better performance across the board, and is relatively insensitive to drive parameters, while the monolithic agent is largely sensitive to the drive parameters, and significantly improves with additional exploration.

### Exploring the intrinsic exploration effect

We investigate further the intrinsic exploration that emerged in the modular models. First, in the simplified case of 2 stats, examined the statistics of the actions selected by each of the two modules. Figure S4 shows that a module could expect to take its preferred action approximately 55% of the time, and that the chance of taking each of the three other (non-preferred) actions ranged approximately between 10% and 20%. This supports our claim that, from the perspective of each module, exploratory (non-preferred) actions were selected approximately randomly in expectation.

Next, we outline one scenario in which this source of exploration was particularly helpful, and may help explain why slow-moving environments were more difficult for the monolithic agent. Occasionally, agents could get stuck in local minima solutions, in which stats steadily decline for long periods of time. One benefit of the non-stationary environment is that a needed resource could randomly appear in front of the agent, helping it solve this problem. An example is shown for the monolithic agent in Figure S5a (top). The purple ‘x’ indicates a resource moving into view, so the agent can begin to replenishing a stat that was previously declining. This helps explain why faster changing environments benefited our agents (i.e., by providing more opportunities to get escape local minima).

As discussed, modular agents also benefit from an intrinsic source of exploration, shown in Figure S5a, (bottom), where the red dots indicate when another module ‘took the wheel,’ helping a module that is stuck find its resource along the way, when its own maximum Q-value was too low to drive the relevant action. Having this additional source of exploration helps explain why the modular agent was less sensitive to the slowly changing resources; Figure S5b shows that a greater proportion of recoveries (defined as a stat declining for at least 200 steps in a row before its resource is found) for the monolithic agent are due to resource location changes, meaning the monolithic agent would be more sensitive to sparsely changing environments (less opportunities for recoveries). Indeed, Figure S5c shows a parametric relationship such that less dependence on resource changes for getting unstuck was associated with better performance for both agents.

Recent work has shown that training a simulated robot with a global reward also decreased its capacity to escape local minima compared with individual limbs learning from local rewards (93). Our results regarding homeostasis, where modularity is with respect to different internal drives, rather than different effectors, indicate that this type of exploration benefit might be a general principle that applies to a variety of important control problems.

### Elaborated results

We show average internal stat trajectories for all our annealing schedules (a more granular look at Figure 3**d**). Figure S6 and Figure S7 show shows these trajectories for a range of number of stats and rates of resource movement respectively.

### Control experiments

Since the modular agent updated each module on each step in the environment, while the monolithic only had a single network to update on each step, a modular agent with *N* modules sampled *N* times more batches from memory overall. One important control experiment therefore involved increasing the number of gradient updates taken per step in the environment for the monolithic model. We show this control in the case of 4 stationary resources, in which the monolithic agent performed 4 gradient update steps (using different sampled batches) at one quarter the learning rate per step in the environment. Our results were unchanged, as seen in Figure S8.

Next, we wondered whether the greatest-mass Q-learning decision process (43) we used was uniqely responsible for our results, or whether other decision processes would display similar results. Therefore, we replaced the decision process in the modular model with the following; the variance over each module’s action values was calculated as *W*_*i*_ = *σ*(*Q*_*i*_(*s, a*)) and the module with the highest W was selected to make the decision on that time-step (122). The idea here was that decisions matter most to the module with the widest range of action values (rather than modules for which all actions mattered about the same amount). The same pattern of results emerged as seen in Figure S9, suggesting that modularity, rather than a particular decision process, was the main driver of our reported effects.

In a similar fashion, we tested whether our results were unique to the particular ϵ-greedy exploration versus some other form of action selection. We therefore repeated simulations in non-stationary and scaled-up environments using SoftMax exploration. That is, actions were selected according to probabilities 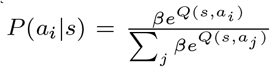, and the inverse temperature parameter *β* was either annealed from 0.2 to 20 on the same schedules as we annealed *ϵ* originally, or held constant at different values. Figure S10 displays these results; the monolithic agent remains strongly exploration-dependent, and struggles in difficult environments compared to the modular agent, further supporting the generality of our main findings.

Next, in order to enhance the opportunity of the monolithic model to make use of attention, we ran a monolithic agent that used a vision transformer rather than an MLP as its backbone. To do this, we used 1x1 patches (i.e. each scalar element of our input vector was passed into the ViT separately to produce a learned embedding, which then were concatenated with a learned positional embedding to be passed to transformer layers). Other than this, we used an out-of-the-box vision transformer (123), trained to output action values and trained identically to our previous models. We used embedding dimension of 64, 3 transformer layers, 6 transformer heads, and output MLP hidden layer dimension of 512 units. This agent still could not outperform the modular agent (Figure S11).

A last note is that the particular reward scaling used, as well as attentional masking, were not crucial for our results in the stationary environment, as a similar pattern of results was observed without masking and whether we used un-normalized rewards, or bounded reward values between -1 and 1 by passing them through a tanh nonlinearity (Figure S12**a** and Figure S12**b**).

### Effect of attentional masking for changing resource locations

Our solution to the increased difficulty of moving resources is outlined in Figure S12. Panels **a** and **b** show a replication of our results with 4 stationary resource locations. Figure S12**c** shows how this pattern of results degrades for 4 moving resources; both modular and monolithic agents have significant difficulty achieving homeostasis. Our hypothesis was that the modular agent could take advantage of its modularity by learning to ignore irrelevant features (i.e. if many resources were moving around, learning to ignore all but the resource it cares about could help it learn). Based on this, we added a learned attentional mask (described in Methods) on each module, as well on the monolithic model for a fair comparison (Figure S12**d**). As predicted, Figure S12**e** strongly enhanced the capabilities of the modular model, and surprisingly also improved the performance of the monolithic model as well (possibly because it added some additional noise into the system given the L1-regularization loss on the learned mask). Figure S12**f** shows the actual learned values of the attentional masks for the modular agent, showing that each module learns to put weights only on the environmental features that are relevant to it.

**Fig. S1.**
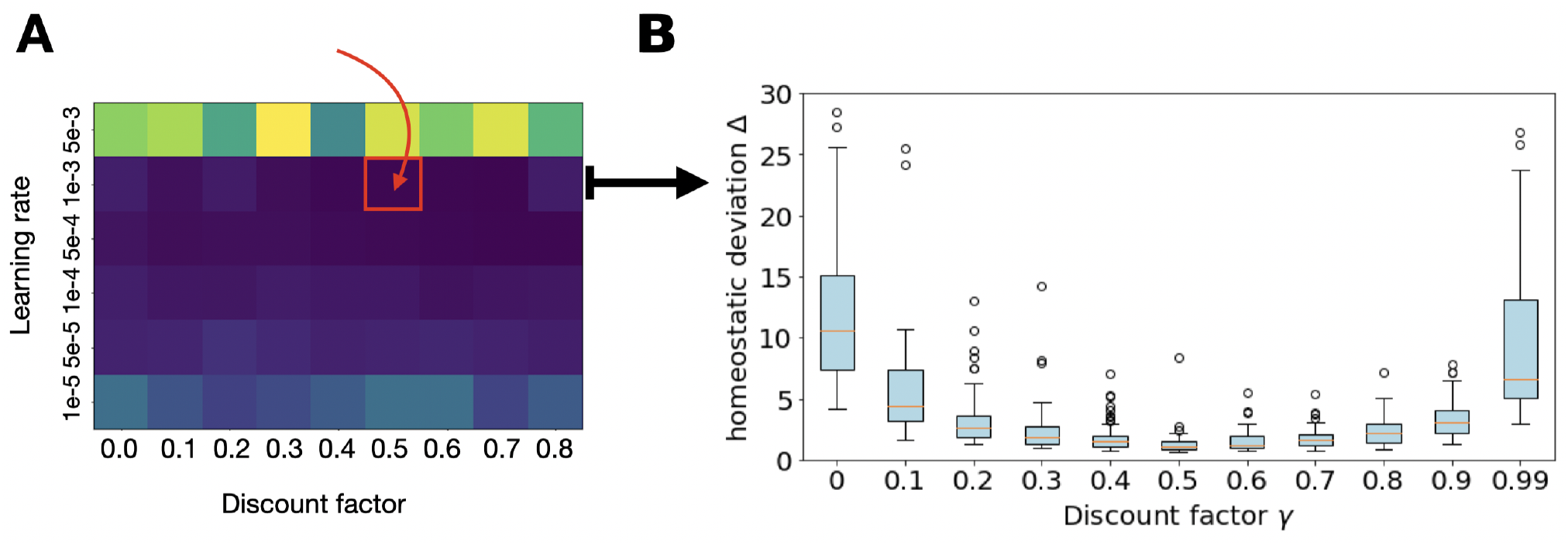
Hyper-parameter search for monolithic model. **a**: Grid-search over learning rates and discount factor. Brightness corresponds to loss; the lowest loss parameter setting of learning rate *α* = 1*e −* 3, discount factor *γ* = 0.5 is indicated by the red square. **b**: A sweep of discount factors at *α* = 1*e −* 3 for *N* = 30 models to provide a sense of performance as a function of this parameter. Boxplots display inter-quartile range and outliers

**Fig. S2.**
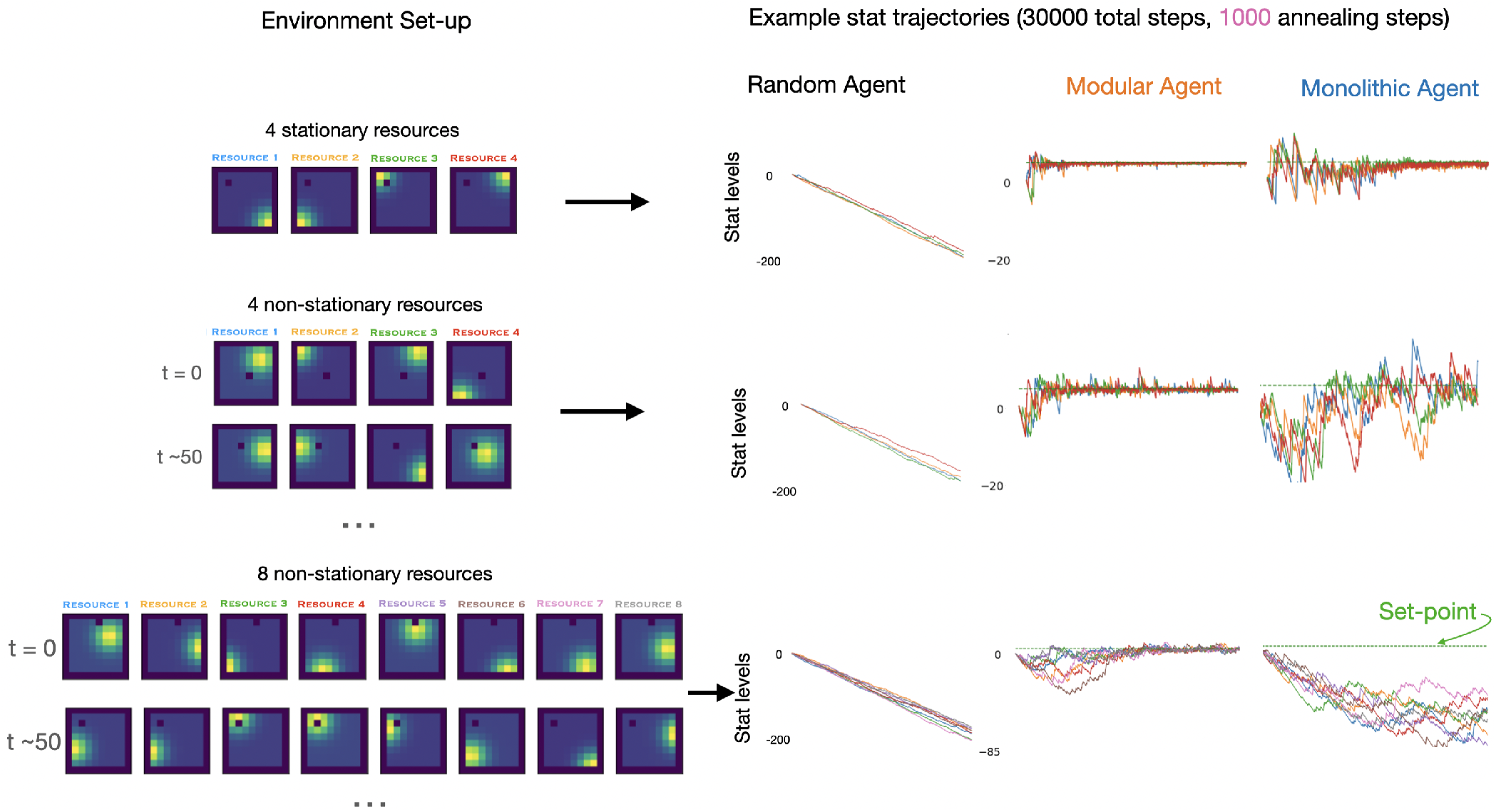
Example learning trajectories for various environment settings. On the left, visual depictions of 3 environment settings are shown; top: 4 resources fixed in 4 corners of grid, middle: 4 resources that eached jumped to a random location according to a poisson process with a rate *λ* = 0.02, bottom: 8 resources that eached jumped to a random location according to a poisson process with a rate *λ* = 0.02. At this rate, resources are expected to change positions every 50 time-steps on average. On the right, an example trajectory of internal stats are shown for 30000 training steps with 1000 epsiolon-annealing steps for a random agent, a modular agent, and a monolithic agent. Separate colours indicate separate internal stats. The set-point of 5 is inidicated by the green dotted line. The modular agent has consistent performance and tends to achieve set-point for all stats. The monolithic agent struggles as task difficulty increases.

**Fig. S3.**
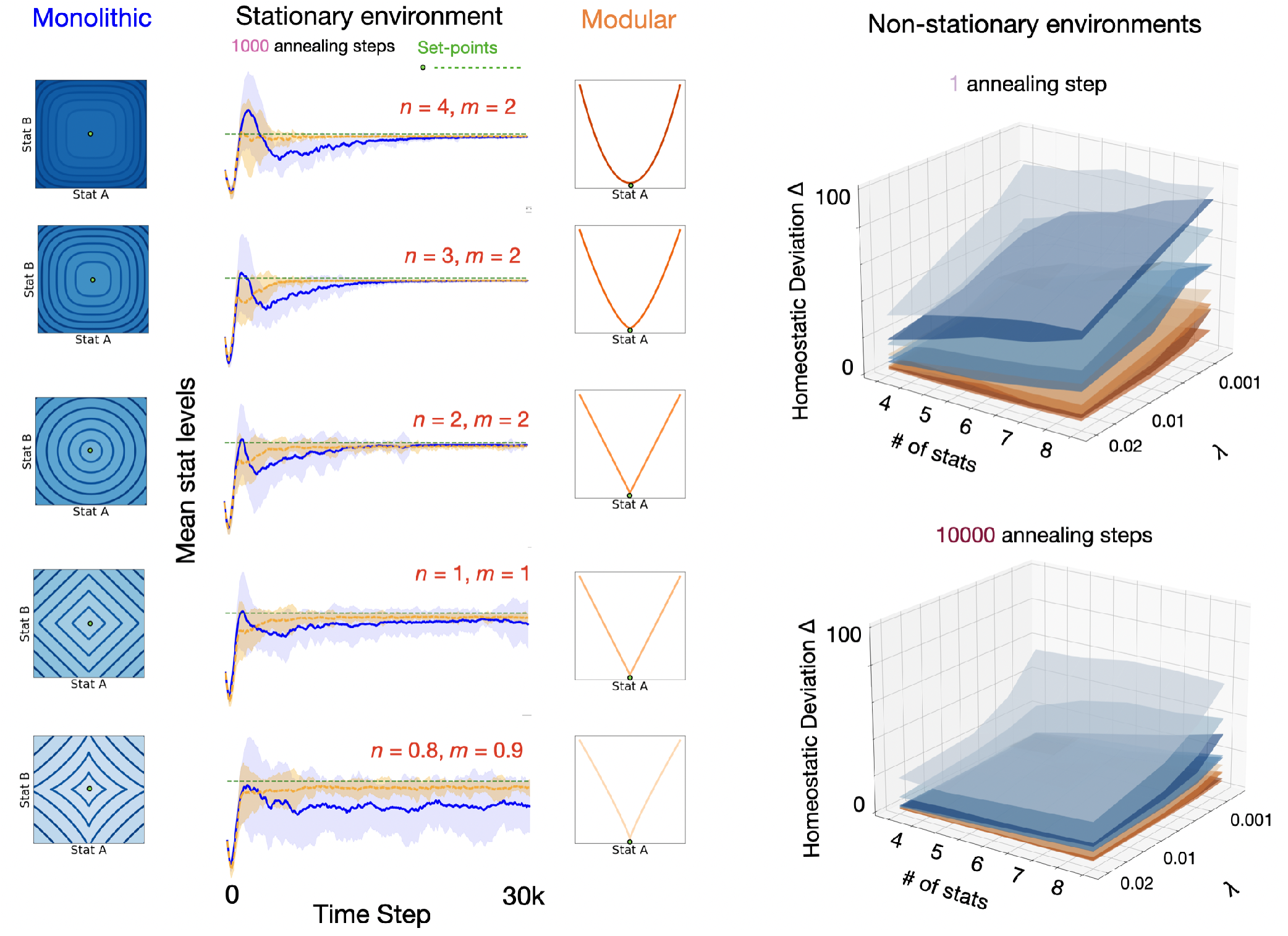
Varying homeostatic drive surface parameters. Left: Replication of Figure 2d (stationary environment, 1000 annealing steps) over additional settings of drive parameters *n* and *m* (red text). Time course of average stat levels during training displayed for monolithic (blue) and modular (orange) agents (N = 30 each; shading shows standard deviation at each time-point, and green dotted line represents set-point of 5 used for all four stats). Corresponding drive surfaces (blue/orange color shading reflecting different (*n, m*) parameters) are displayed for reference (2D for monolithic, shown to the left of the plots; and 1D for modular, shown to the right). Right: Replication of low and high exploration conditions from Figure 3d (non-stationary environment) over additional settings of drive parameters *n* and *m*, averaged over N=60 agents (blue/orange color shading reflecting the same (*n, m*) parameters as the corresponding drive surface schematics on the left). The left and right horizontal axes of each plot represent number of stats and rate of resource location change (*λ*) respectively, and the vertical axis displays homeostatic performance. The modular agent (orange surfaces) exhibit superior performance across all conditions and parameter settings.

**Fig. S4.**
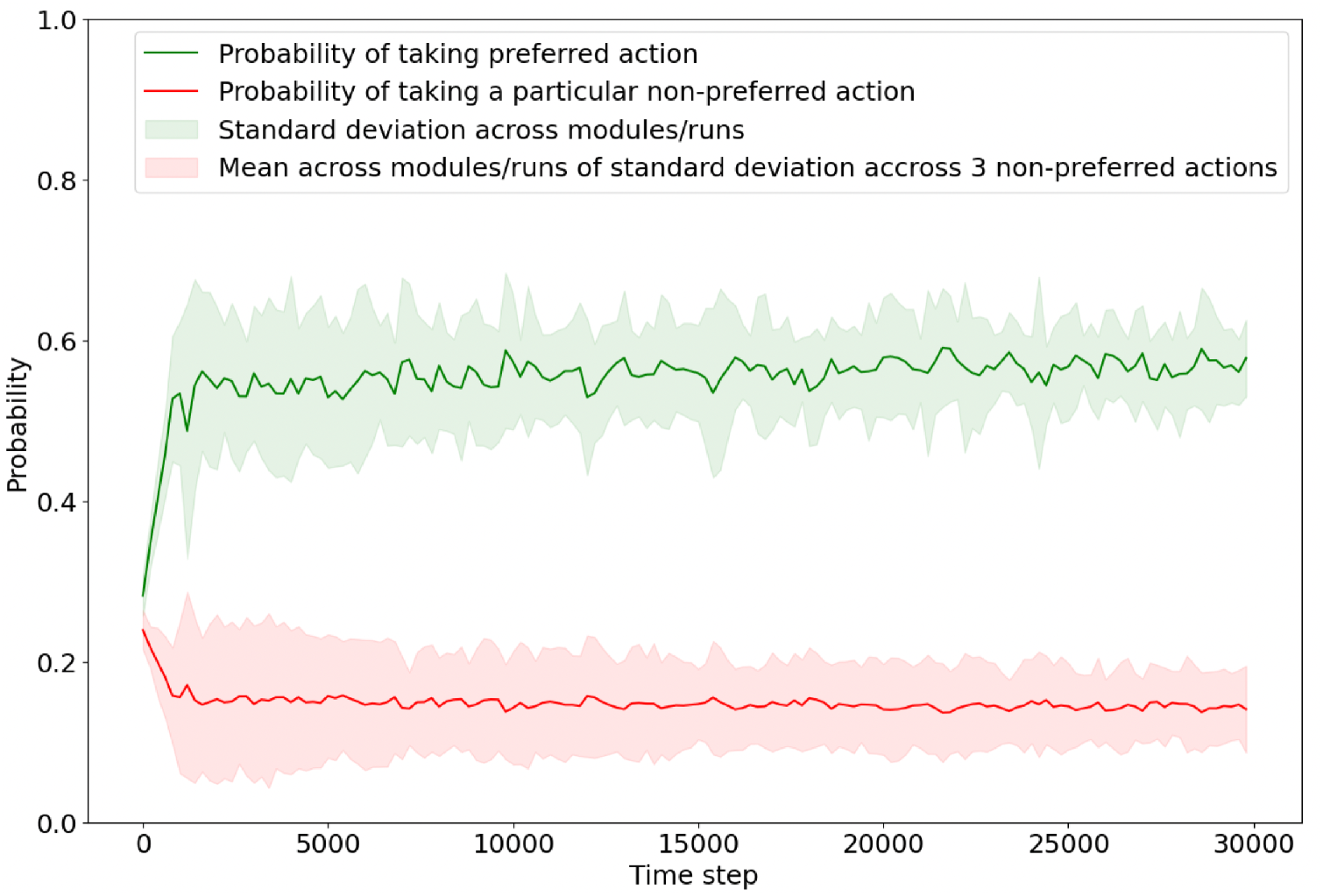
Quantifying action probabilities in the modular agent. We used a simplified non-stationary environment (2 stats, 1000 annealing steps, *λ* = 0.01) to calculate the probability of a module taking its preferred action (green) or any particular non-preferred action (red). The red shading represents the average standard deviation across non-preferred actions, showing that non-preferred actions were selected approximately randomly in expectation (i.e. they all had a similar probability of being selected, conditioned on the preferred action of the agent).

**Fig. S5.**
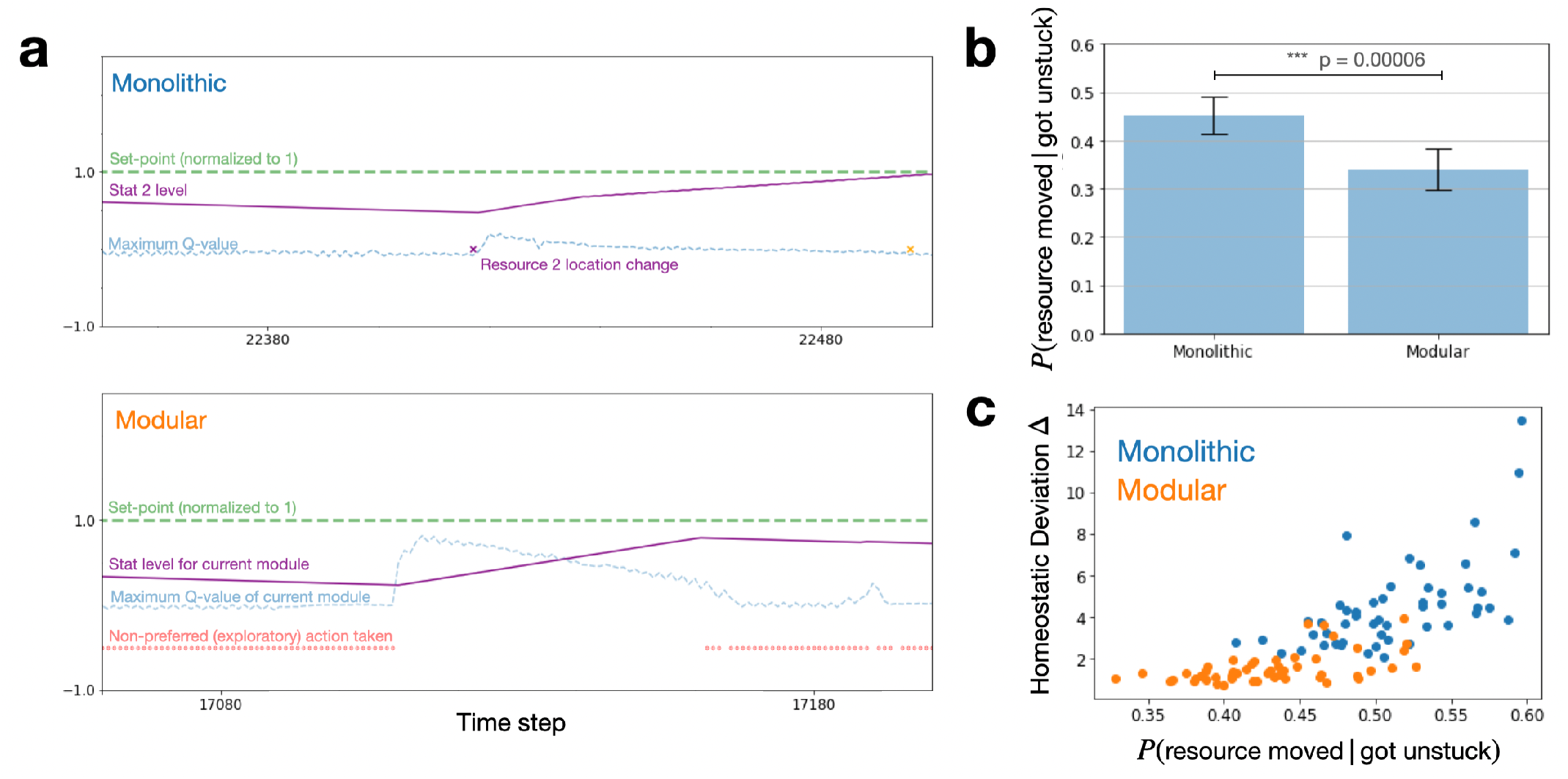
Escaping local minima in changing environments. **a**: Top: Example scenario in which a resource location change (purple x) helps a monolithic agent replenish stat 2 (purple line), that was continually depleting due to an unchanging maximum Q-value (blue dotted line). Bottom: An analogous scenario for a modular agent, but here, non-preferred actions taken by a different module (red dots) help the agent find the resource it needs. **b**: The probability that a resource moved location within 10 steps before an agent ‘got unstuck’ (defined as a particular stat being replenished after a period of at least 200 continuous steps of decline). Error bars reflect mean and standard deviation for N=60 agents, p-value calculated with two-sample t-test. The modular agent was more likely to have got unstuck without any preceding resource location. **c**: The probabilities from panel **b** correlated with homeostatic performance for both agents; the more an agent relied on resource changes to escape local minima, the worse the overall performance.

**Fig. S6.**
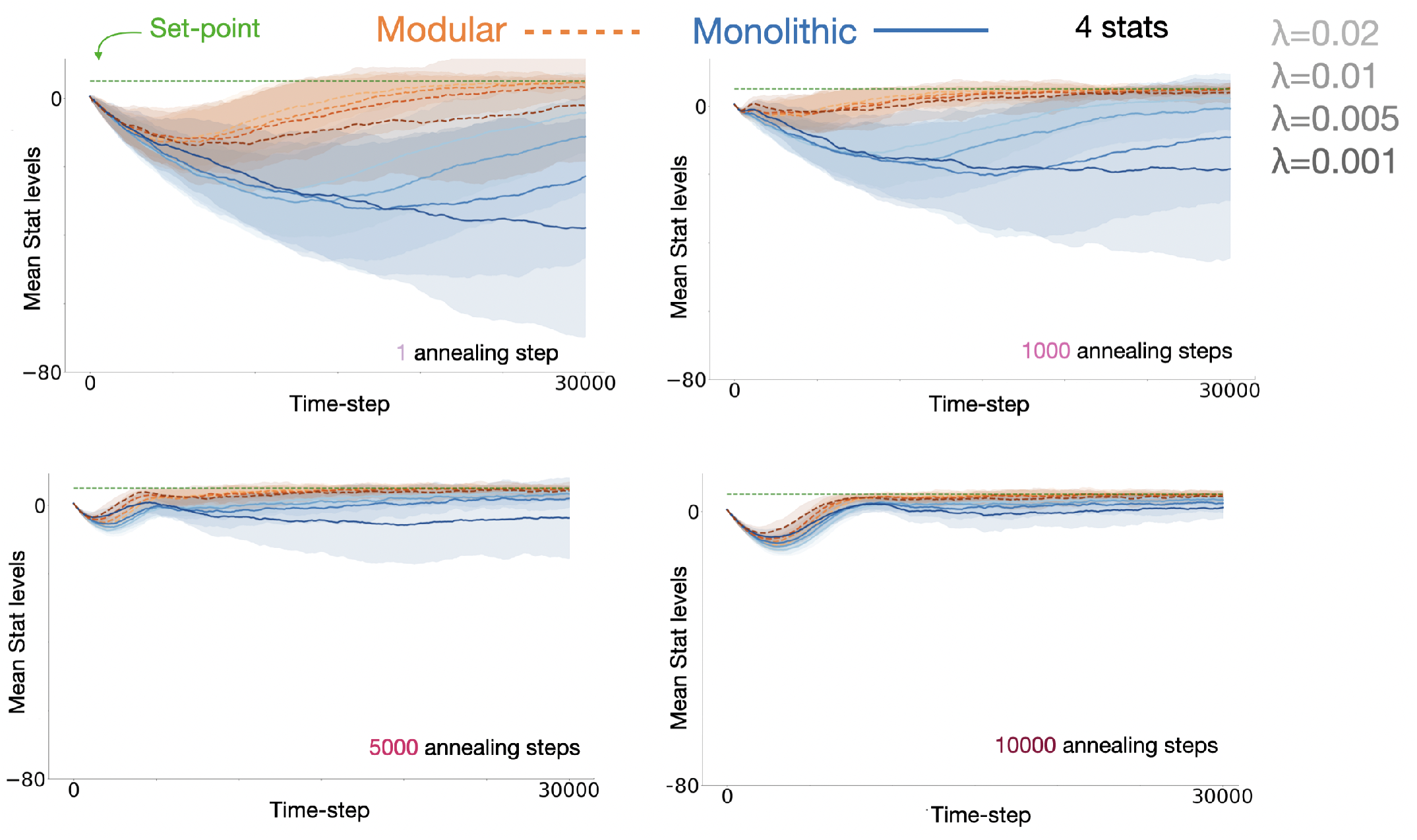
Modular (orange) and monolithic (blue) internal stat trajectories for different amounts of epsilon annealing, with 4 internal stats. Darker colors indicate decreasing Poisson rate of resource location changing (i.e. resources move locations more slowly). Homeostasis is worse for both agents at slower change rates, but the effect is significantly greater for the monolithic agent - the worst case modular agent is better than the best-case monolithic agent in all cases. Shading represents standard deviation across N=100 models.

**Table S1.** Parameter settings for environment and models assuming 4 resources

| Model | Monolithic | Modular | Environment |  |
| --- | --- | --- | --- | --- |
| Trainable parameters | 1.09e6 | 1.09e6 | # of resources $N$ | 4 |
| MLP hidden layer units | 1024 | 500 | HRRL exponents $(n, m)$ | (4, 2) |
| Learning rate | 1e-3 | 1e-3 | Stat set-points $h_i^*$ | 5 |
| Discount factor $\gamma$ | 0.5 | 0.5 | Initial stat levels $h_{i,t=0}$ | 0 |
| Memory buffer capacity | 30k | 30k | Resource locations $\mu_x, \mu_y$ | variable |
| Target network update frequency | 200 | 200 | Resource covariance $\Sigma$ | $\begin{pmatrix} 1 & 0 \\ 0 & 1 \end{pmatrix}$ |
| Batch size | 512 | 512 | Stat depletion per step | 0.01 |
| Initial $\epsilon$ | 1 | 1 | $R_{\text{thresh}}$ | 95% |
| Final $\epsilon$ | 0.01 | 0.01 | mask sparsity weight $\beta$ | 0.0001 |

**Fig. S7.**
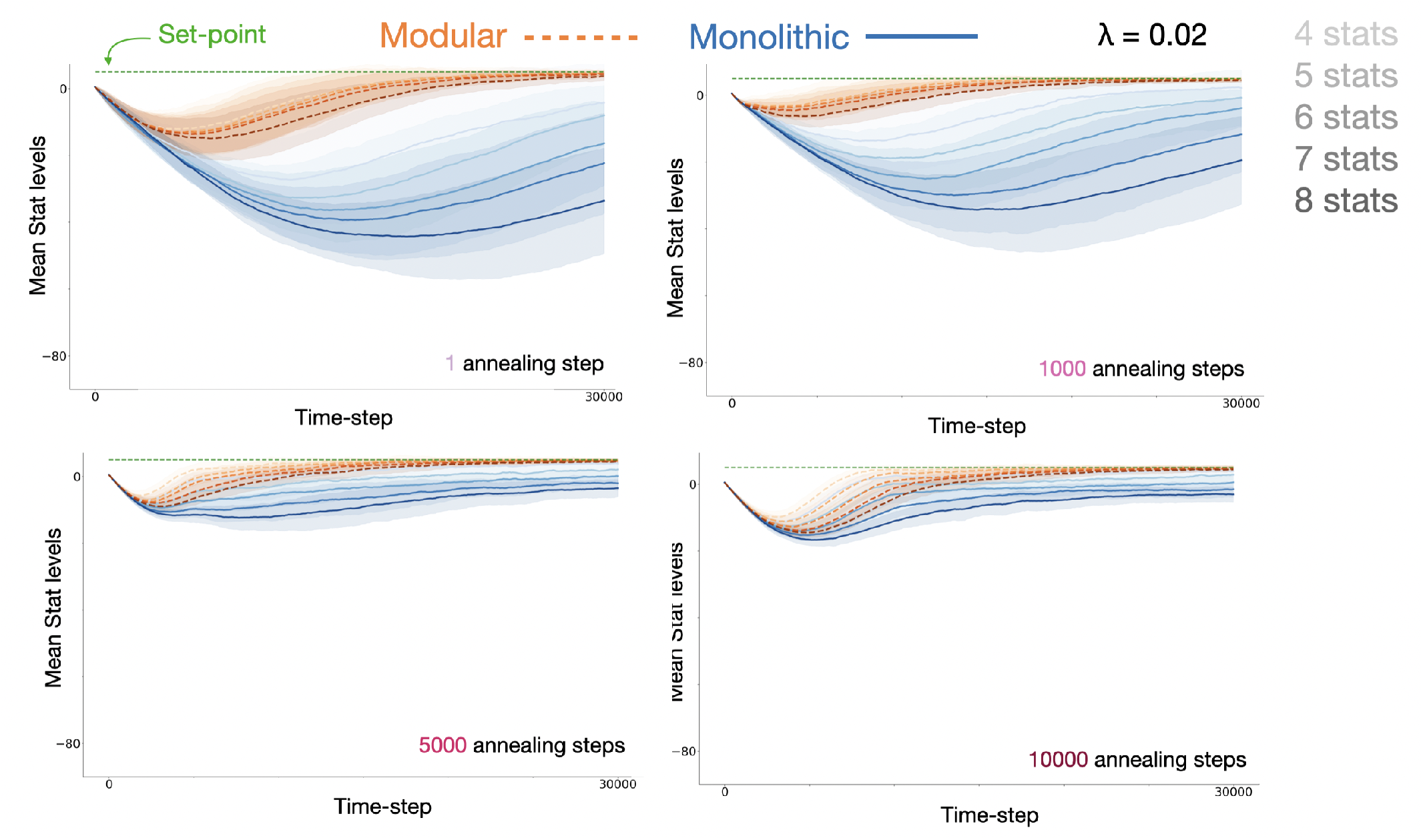
Modular (orange) and monolithic (blue) internal stat trajectories for different amounts of epsilon annealing, resource locations changing at Poisson rate of *λ* = 0.02. Darker colours indicate increasing number of needs. Homeostasis is worse for both agents with more internal stats, but the effect is significantly greater for the monolithic agent - the worst case modular agent is better than the best-case monolithic agent in all cases. Shading represents standard deviation accross N=100 models.

**Fig. S8.**
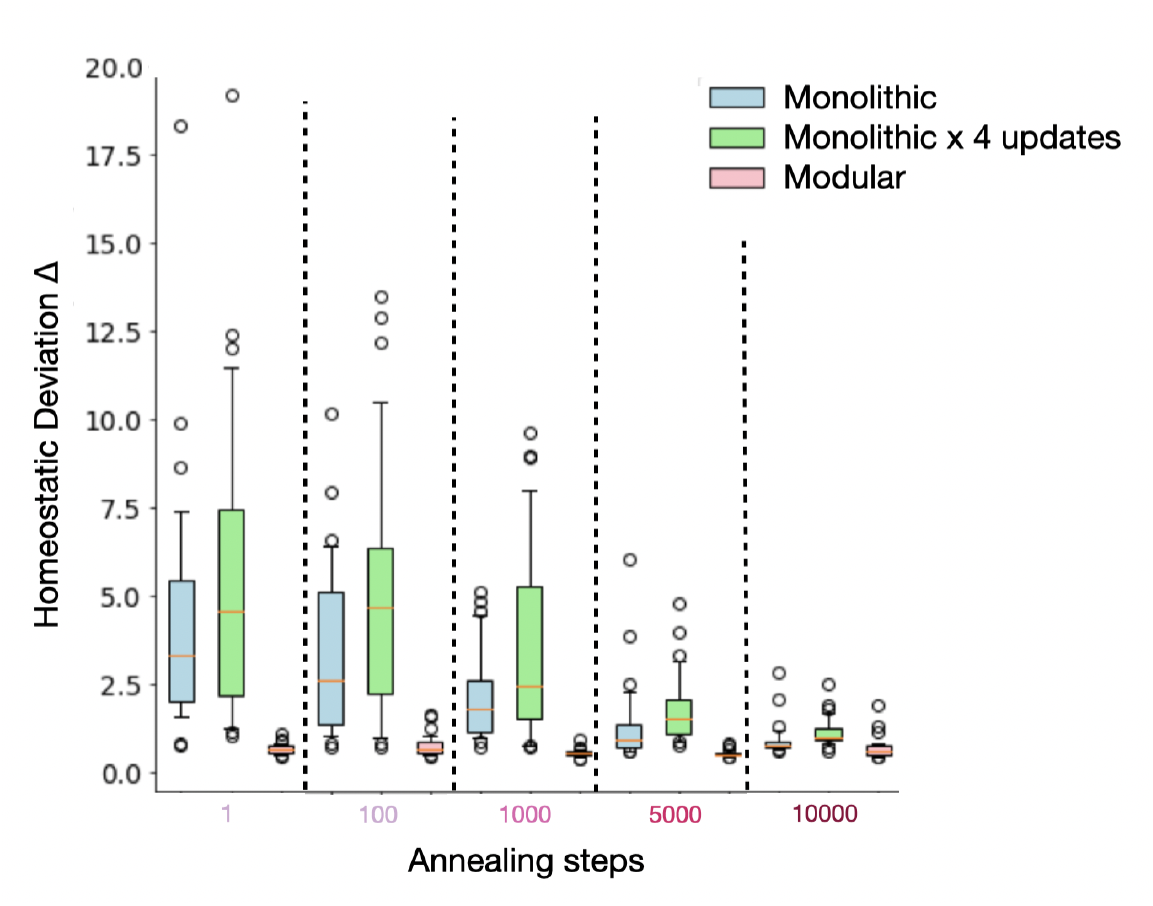
Comparison of modular (pink), monolithic (light blue) and monolithic with 4 gradient updates per step (green) models in a stationary environment where resources are fixed in the four corners of the grid-world. Homeostatic performance is calculated as in Eq. [8] (lower is better). The modular agent is relatively unaffected by the exploration annealing schedule, whereas both monolithic models require significant exploration to have comparable performance. Boxplots display inter-quartile range and outliers for N=30 models.

**Fig. S9.**
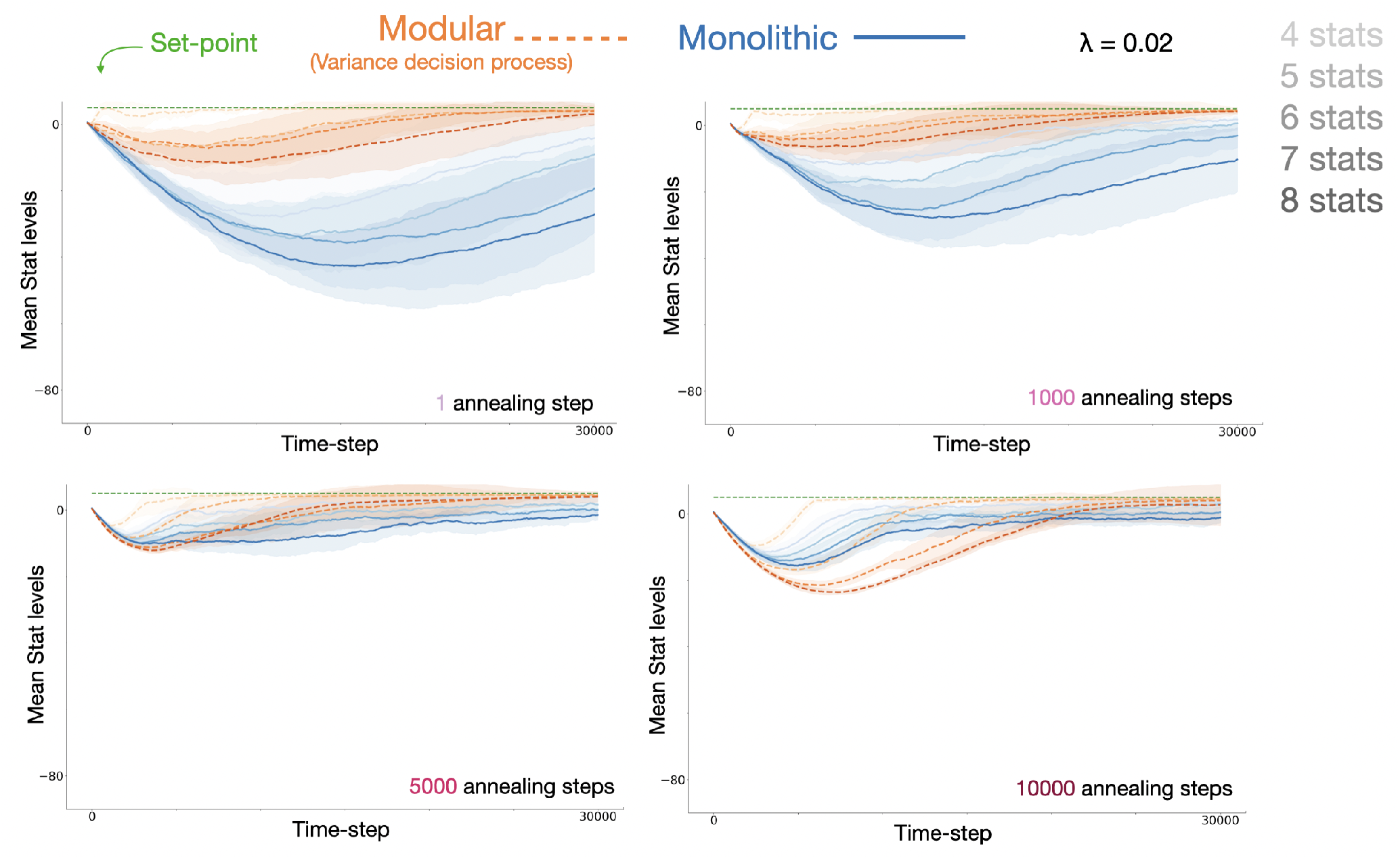
Modular agent using the variance decision process (orange) and monolithic (blue) internal stat trajectories for different amounts of epsilon annealing, resource locations changing at Poisson rate of *λ* = 0.02. Darker colours indicate increasing number of needs. Homeostasis is worse for both agents with more internal stats, but the effect is again significantly greater for the monolithic agent - the worst case modular agent is better than the best-case monolithic agent in all cases. Shading represents standard deviation across N=30 models.

**Fig. S10.**
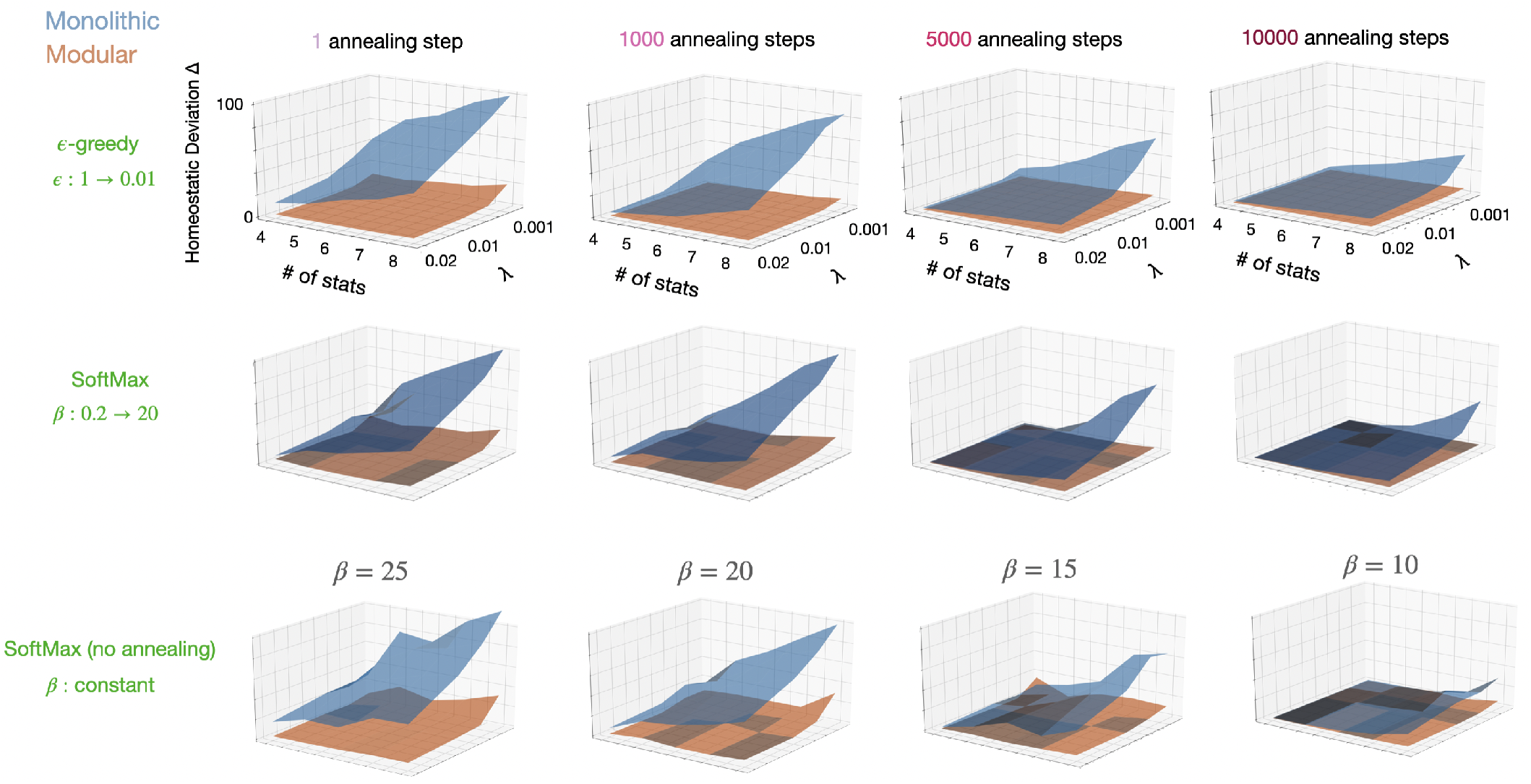
Replication of results using SoftMax exploration. Top row: original results in non-stationary environment (from Fig. 3d). The left and right horizontal axes of each plot represent number of stats and rate of resource location change (*λ*) respectively, and the vertical axis displays homeostatic performance. Subsequent rows share axis labels. Middle row: Corresponding results using SoftMax exploration with inverse temperature *β* annealed from 0.2 to 20 on the same schedules as originally used for *ϵ*-greedy. Bottom row: Results using SoftMax exploration but with constant *β* throughout training of 25, 20, 15, or 10.

**Fig. S11.**
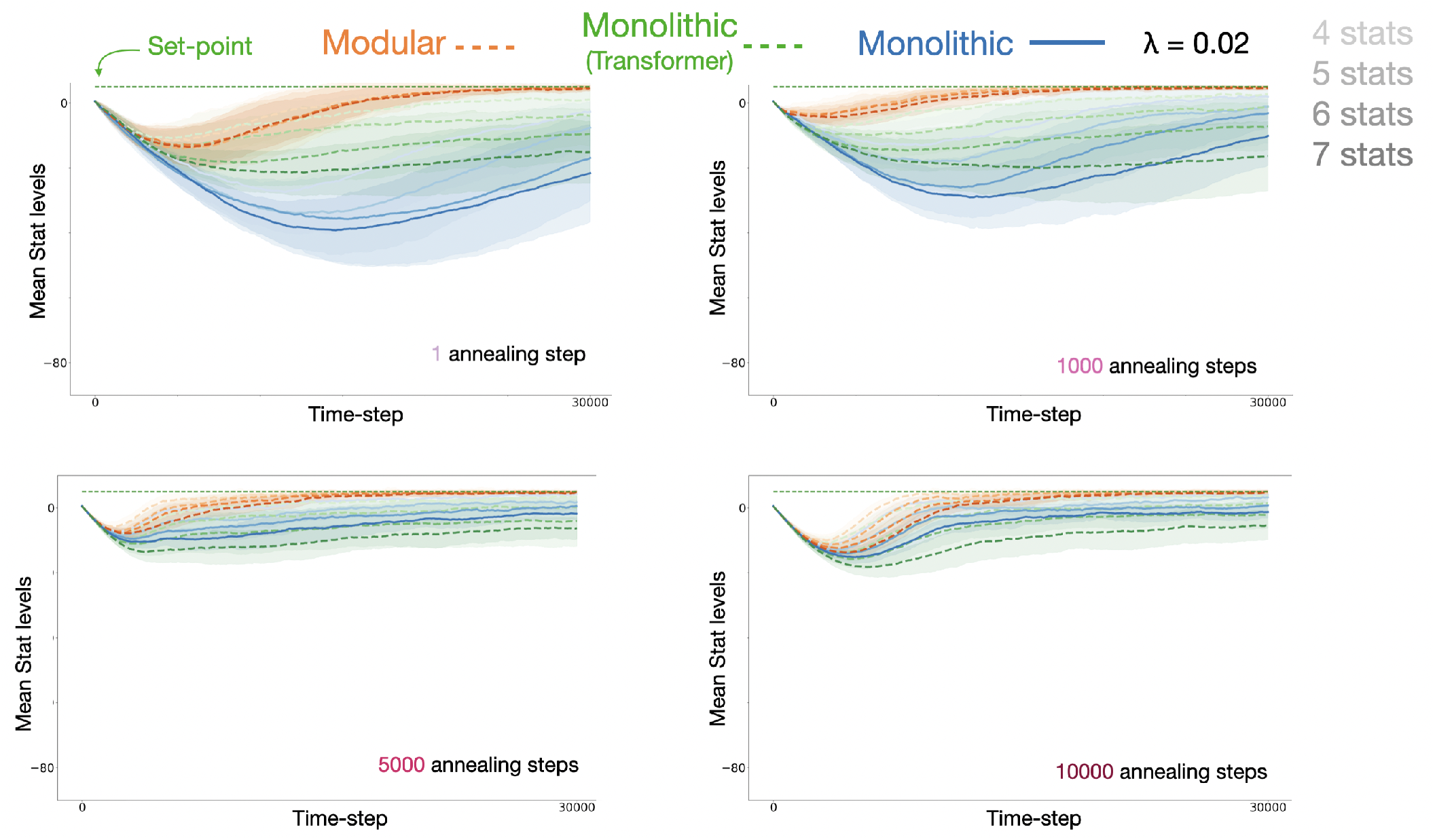
Modular agent (orange), monolithic agent (blue) and monolithic agent using vision transformer (green) internal stat trajectories for different amounts of epsilon annealing, resource locations changing at Poisson rate of *λ* = 0.02. Darker colours indicate increasing number of needs. The modular agent is still the best performing model. Shading represents standard deviation across N=30 models.

**Fig. S12.**
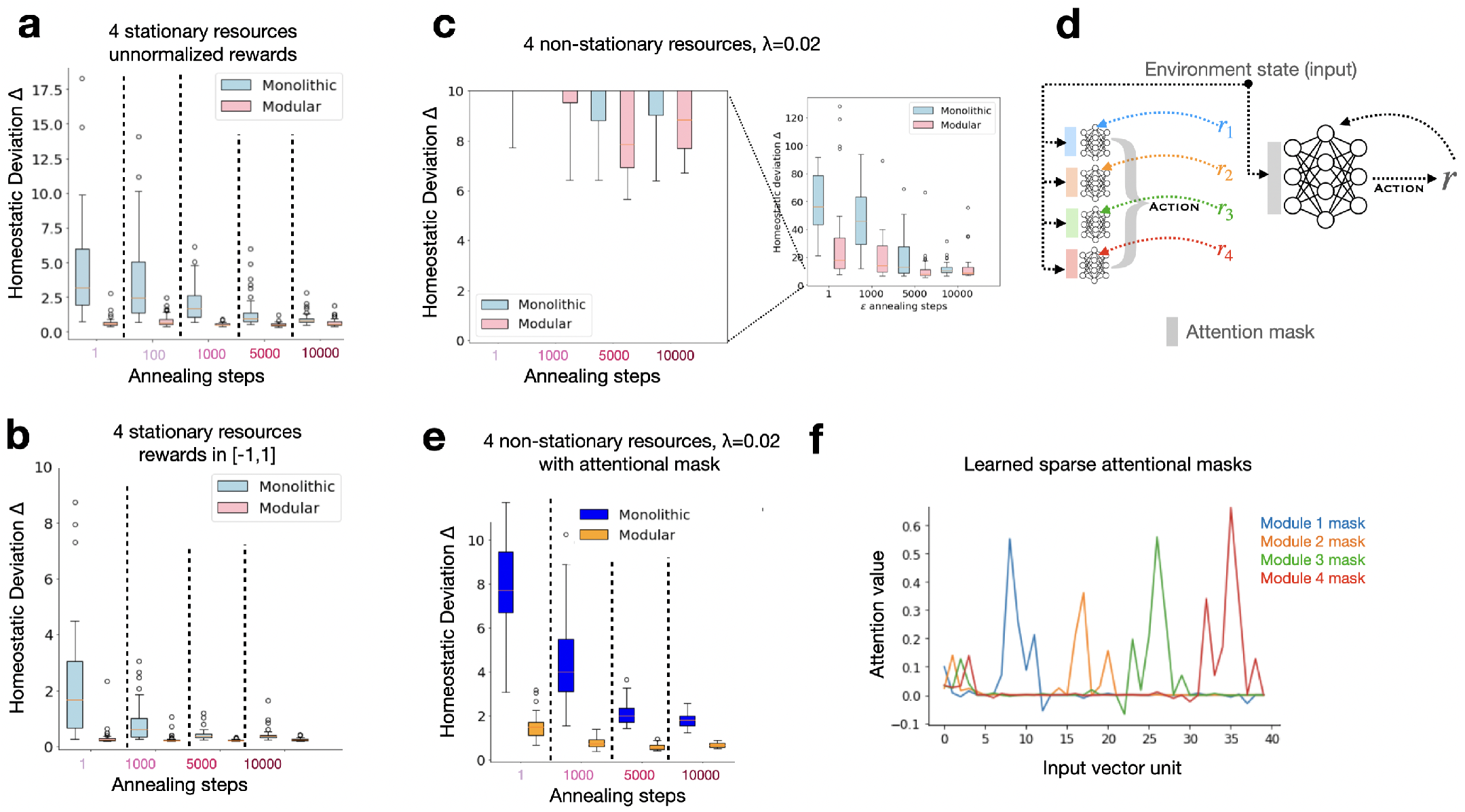
Effect of learning an attentional mask. Lightblue/pink color schemes are used for results without attention, blue/orange scheme indicates the addition of attention. **a**: replication of results in stationary environment using unnormalized rewards (N=30). **b**: Original stationary environment results reported in main text. **c**: Using the same settings as in **b**, the degraded performance of monolithic and modular models for non-stationary resources locations. Inset shows the complete y-axis scale. **d**: Learned attention masks were vectors the same size as the state input vectors, that were element-wise multiplied with the inputs before they were passed into the DQN networks. **e**: Results as reported in the main text with attention masking. **f**: Learned values of the 40-element attention masks for the modular agent after training. The first 4 elements of the input vector are 4 internal stats, and the following 4 sets of 9 elements are the 3x3 egocentric views of the agent of each resource map in order. It can be seen that the attention values for module 1, for example, are highest on internal stat 1 and on resource map 1, and fall to 0 elsewhere.

